# Molecular regulation of GPCR-G-protein-governed PIP3 generation and its adaptation

**DOI:** 10.1101/2022.08.31.506078

**Authors:** Dhanushan Wijayaratna, Kasun Ratnayake, Sithurandi Ubeysinghe, Dinesh Kankanamge, Mithila Tennakoon, Ajith Karunarathne

## Abstract

Phosphatidylinositol (3,4,5) trisphosphate (PIP3) is a plasma membrane-bound signaling phospholipid involved in many cellular signaling pathways that control crucial cellular processes and behaviors, including cytoskeleton remodeling, metabolism, chemotaxis, and apoptosis. Therefore, defective PIP3 signaling is implicated in various disease driving processes, including cancer metastasis, diabetes, obesity, and cardiovascular diseases. Upon activation by G protein-coupled receptors (GPCRs) or receptor tyrosine kinases (RTKs), phosphoinositide-3-kinases (PI3Ks) phosphorylate phosphatidylinositol (4,5) bisphosphate (PIP2), generating PIP3. Interestingly, though the mechanisms are unclear, PIP3 produced upon GPCR activation attenuates within minutes, indicating a tight temporal regulation. Our data show the subcellular redistributions of G proteins govern this PIP3 attenuation in the presence of sustained receptor stimulation, and thus meet the definition of signaling adaptation. Interestingly the observed adaptation of PIP3 was Gγ subtype-dependent. Considering distinct cell-tissue-specific Gγ expression profiles, our findings not only demonstrate how the GPCR-induced PIP3 response is adapted but also show how diversely this adaptation process is regulated by the dominant Gγs of a cell.

## 1. Introduction

The cell is the ultimate building block of living organisms. The conditions of the cell exterior regulate cellular functions^1^. These extracellular conditions are perceived by the cell through stimuli. Stimuli are chemical or physical signals that influence the homeostasis of a cell. Typical stimuli include hormones, neurotransmitters, and odorants^2^. Receptors on the cell surface sense numerous stimuli and initiate signaling at the inner leaflet of the plasma membrane, which then broadcasts to subcellular domain, controlling a variety of physiological processes^2,3^. Interestingly, cells manage to desensitize itself to sustained stimulation, protecting the cell from excessive and deleterious signaling^3^, a phenomenon known as ‘adaptation.’ Adapting biological systems are found in species ranging from bacteria to eukaryotes. In eukaryotes, signaling adaptation is involved with many crucial physiological processes, including pain, vision, and olfaction^3,4^. A variety of mechanisms have been identified thus far, which cope with sustained stimulation that allows cells to adapt. For instance, sensory neurons show an immediate response to a stimulus, however, with time, the response level will decrease to the prestimulus level despite continuous stimuli. Only another stronger stimulus will be able to elicit a response in this adapted system^3^.

Literature provides evidence for adapting signaling responses mediated by GPCRs and G proteins^2,3,5^. Ras activation, phosphoinositide production, Protein kinase B (PKB) activation, cAMP production, and calcium influx are key GPCR-mediated signaling processes that exhibit adaptation upon continuous stimuli^3^. For instance, the adaptation of PIP2 signaling upon continuous stimulus has been shown previously^5^. As major regulators of cell fate and pathogenesis, kinases and phosphatases, play crucial roles in phosphoinositide signaling by regulating cellular phosphoinositide levels^6^. For example, dysregulated expression and compartmentalization of specific kinases in the proper balance of phosphorylation-dephosphorylation events in cells have been implicated in cancer^6^. By catalyzing the phosphorylation of PIP2, Phosphoinositide 3-kinases (PI3Ks) generates PIP3, which is a crucial signaling phospholipid^7-9^. So far, three classes of PI3Ks (class I, II, and III) have been identified^9^. Though there are three classes of PI3Ks with multiple isoforms in each class^10^, only class I_A_ PI3Kβ (p110β catalytic and p85 regulatory) and I_B_ PI3Kγ (p110γ catalytic and p101 regulatory) are activated by Gβγ released upon GPCR stimulation^11,12^. These catalytic-regulatory dimers are cytosolic and recruited to the plasma membrane upon Gβγ generation^12-14^. PIP3 is involved in crucial cellular regulation processes such as phagocytosis, localization of protein kinases, regulation of GTPases^15^, and multiple physiological processes, including cell migration, tissue regeneration, and chemotaxis^16^. In addition to GPCRs, PIP3 also are generated upon activation of other signaling pathways, including receptor tyrosine kinases such as insulin receptor (InsR)^17^. Phospholipids act as docking sites for intracellular signaling proteins by recruiting multiple effector proteins to the plasma membrane. PIP3 recruits effectors to the plasma membrane mainly through the interactions with Pleckstrin homology (PH) domains. PI3K-induced PIP3 generation leads to Akt and mTOR pathway activation, reducing apoptosis and enhancing cell proliferation^18^. Hence, many viral GPCRs activate PI3Ks to produce PIP3, benefiting the viral life cycle. Furthermore, viral GPCRs are constitutively active, and thus PI3Ks could remain active indefinitely^19^. Therefore, PI3K/Akt/mTOR signaling becomes tumorigenic and allows viral genome replication^20^.

PI3K/Akt pathway is tightly regulated in healthy cells^7^. Previously, it has been shown that GPCR activation-induced PIP3 exhibits adaptation within minutes upon a continuously applied stimulus^21-25^. Different models of cellular chemotaxis signaling have been proposed to describe a mechanism for the chemotaxis adaptation process^21,22^. One of the most popular mechanisms explaining bacterial chemotaxis is the ‘Barkai-Leibler’ model^26^, which describes a mechanism for chemotaxis adaptation in a bacterial cell. When a chemotactic ligand binds the receptor, it induces flagella movement due to the kinase activity of CheA protein. However, CheA also phosphorylates CheB. Thus, phosphorylated CheB demethylates the receptor, which causes adaptation of chemotaxis. Methylation of the receptor is essential for normal chemotaxis in bacteria^27^. However, an explanation at the molecular level for PIP3 adaptation in eukaryotic cells is lacking. The ‘local effector global inhibitor’ (LEGI) model shows that *Dictyostelium discoideum* cells form localized PIP3 in response to a chemoattractant gradient^21^. Distribution of the effector (PI3K) and inhibitor (PTEN) was suggested as underlying mechanisms. However, when GPCRs across a cell are activated asymmetrically, a localized adaptation-resistant PIP3 is formed^20^. The LEGI model does not explain how PI3K or PTEN is regulated to distribute unevenly across a cell to induce localized PIP3. Since the literature does not provide much information on the underlying molecular mechanisms of PIP3 adaptation, we wanted to explore the fundamentals behind this adaptation process.

Here, using live-cell imaging, subcellular optogenetic GPCR, G protein activation, and single-cell analysis utilizing multiple biosensors, we examined PIP3 dynamics in live cells. Here, we show that PIP3 depletes on the plasma membrane in the presence of continuous stimuli, indicating it agrees with the definition of adaptation. Additionally, we observed distinct PIP3 regulation in GPCR-G protein activity compared to RTK activity. Our results show the molecular reasoning behind the distinct regulatory mechanisms in GPCR-G protein activation-mediated PIP3 generation and adaptation. The molecular underpinning of the PIP3 adaptation process demonstrates how a vital cellular signaling machinery is implicated in many essential physiological functions and pathophysiology.

## 2. Results and Discussion

### 2.1 Gi/o-coupled GPCRs induce a self-attenuating PIP3 production

Gi/o-coupled GPCR activation stimulates PI3Kγ at the plasma membrane leading to PIP3 generation^14,15,28^. We activated Gi/o-coupled alpha 2 adrenergic receptors (α2AR) using 100 μM norepinephrine and examined the PIP3 generation in RAW264.7 mouse macrophage cells. PIP3 response was observed using the PIP3 sensor, Akt-PH-Venus, which is cytosolic in the absence of PIP3 and translocates to the plasma membrane upon PIP3 generation. We observed a robust PIP3 production with a half-time (t_½_) of = 109.78 ± 4.01 s (Fig. 1A, up to 300 s, and plots). Here, we measured PIP3 production using the reduction in cytosolic PIP3 sensor fluorescence due to its plasma membrane recruitment to avoid experimental artifacts because PIP3 causes cell changes, introducing artifacts to membrane fluorescence measurement. Interestingly, after PIP3 production reached the maximum, Akt-PH returned to the cytosol with a t_½_of = 433.85455 ± 12.13 s, indicating the reduction of PIP3 at the plasma membrane (Fig. 1A, 900 and 1500 s and plots). Since we did not remove the α2AR agonist, norepinephrine, if GPCRs and G proteins stayed active during this period, the observed significant PIP3 reduction meets the definition of signaling adaptation^1^.

**Figure 1.**
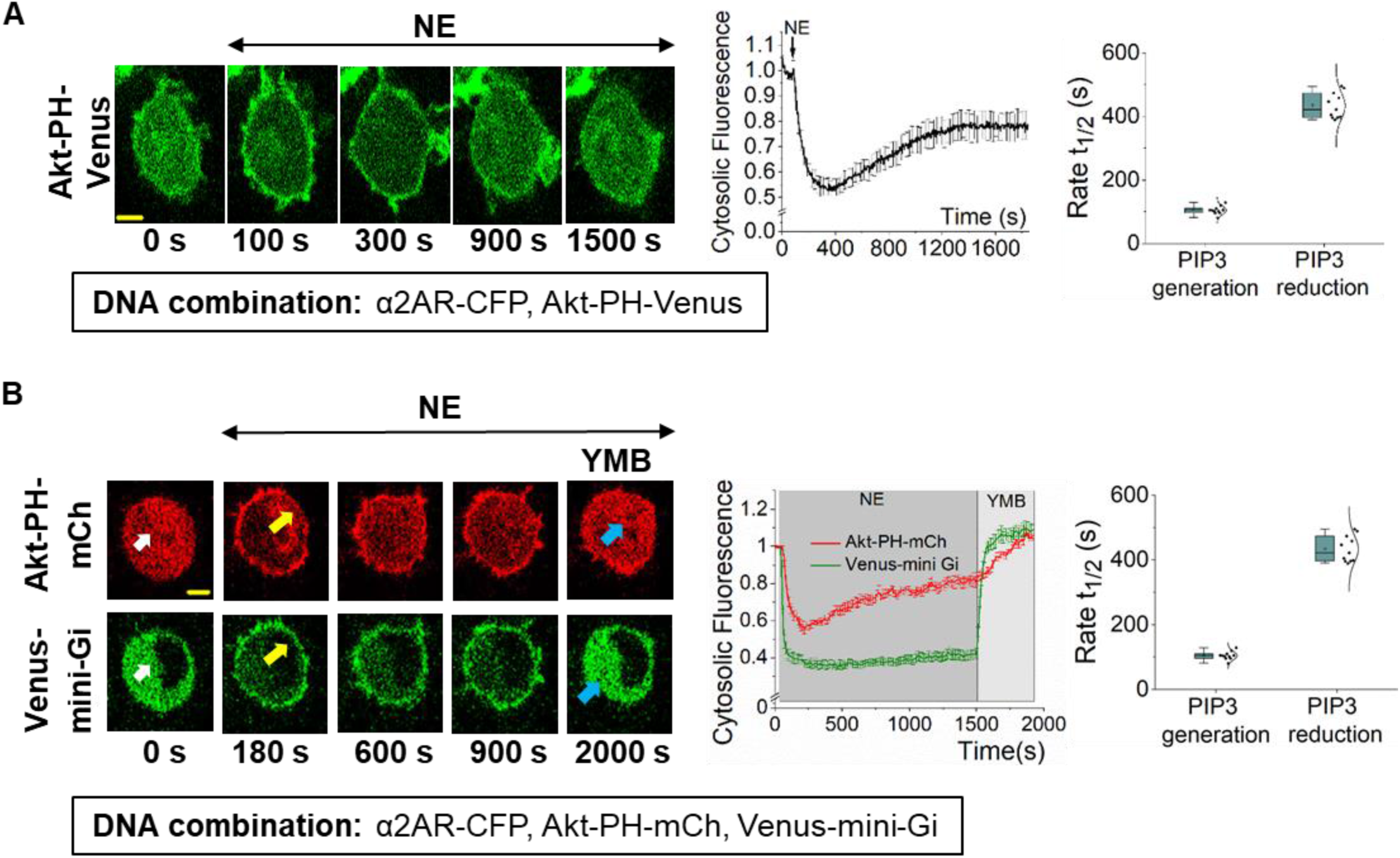
While GPCR and G proteins stay active, Gi/o-GPCR induced PIP3 subsides. **(A)** RAW264.7 cells exhibited robust PIP3 production upon α2AR activation, which shows subsequent significant attenuation. Images: RAW264.7 cells expressing Akt-PH-Venus (PIP3 sensor) and α2AR-CFP (not shown). Confocal time-lapse imaging of PIP3 sensor dynamics was performed using 515 nm excitation. α2AR was stimulated using 100 μM NE at 1 minute. Images show PIP3 sensor translocation from the cytosol to the PM upon α2AR activation, which peaks at 300 s, and significantly reverses back to the cytosol in 5-10 minutes. The corresponding plot shows PIP3 sensor dynamics in the cytosol of the cells. The whisker box plot shows the half-time (t_½_) of PIP3 generation and adaptation (n = 13). **(B)** RAW264.7 cells expressing with α2AR-CFP, Akt-PH-mCh, and Venus-mini-Gi exhibited simultaneous mini-Gi PM-recruitment and PIP3 production (yellow arrows) upon α2AR activation. 100 μM NE was added at 1 minute to activate α2AR. Mini-Gi stayed recruited to the PM, while PIP3 showed adaptation. Upon addition of α2AR inhibitor yohimbine (50 μM), both mini-Gi and PIP3 sensors showed complete reversal to the cytosol (blue arrows). Cells were imaged using 515 nm (to monitor mini-Gi), and 594 nm (to capture the PIP3 sensor) excitation. The corresponding plot shows the mini-Gi (green) and PIP3 (red) sensor dynamics in the cytosol of the cells. The whisker box plot shows the half-time (t_½_) of PIP3 generation and adaptation rates (n = 11). Average curves were plotted using cells from ≥ 3 independent experiments. ‘n’ denotes the number of cells. Error bars represent SEM (standard error). The scale bar = 5 μm. GPCR: G protein-coupled receptor; PIP3: Phosphatidylinositol 3,4,5 triphosphate; α2AR: Alpha-2-adrenergic receptor; NE: Norepinephrine; CFP: Cyan Fluorescence protein; mCh: mCherry; PM: Plasma membrane

GPCR-G protein activation-induced cellular processes are usually terminated upon GPCR deactivation due to agonist removal, exposing the receptor to antagonists or GPCR desensitization and internalization upon their phosphorylation^29,30^. Therefore, using a fluorescently-tagged mini-G protein, we examined whether a reduction of cell surface concentration of active α2AR is responsible for the observed PIP3 reduction. Mini-G proteins are cytosolic sensors recruited to active GPCRs upon agonist addition^31^. Since α2AR is Gi/o-coupled, we expressed Venus-mini-Gi and Akt-PH-mCherry (Akt-PH-mCh) to monitor the status of the receptor and PIP3 response simultaneously in the same cell. Before activation, both sensors were cytosolic (Fig. 1B, 0 min-white arrows). Norepinephrine addition induced simultaneous PIP3 generation and mini-Gi recruitment to the plasma membrane (Fig. 1B- yellow arrows). Continuous monitoring showed that PIP3 gradually and significantly disappeared with a t_½_of = 426.39 ± 19.54 s (Fig. 1B-red plot). However, mini-Gi stayed on the plasma membrane, indicating active-state GPCR (Fig. 1B- green plot). Therefore, the observed PIP3 disappearance meets the criteria of a partial adaptation because, during the process, GPCRs remained active. The addition of α2AR antagonist, yohimbine (50 μM) deactivated α2ARs, resulting in a complete reverse translocation of mini-Gi and the PIP3 sensor to the cytosol (Fig. 1B- blue arrows). These data collectively show that the diminished GPCR activity is not a significant factor in the observed partial adaptation of PIP3.

### 2.2 PIP3 adaptation is not a result of the substrate, PIP2 or product, PIP3 depletion

Active PLCβ hydrolyzes PIP2, generating inositol triphosphate (IP3) and diacylglycerol (DAG), and PIP2 is the substrate for PI3K to generate PIP3^32^. Though GαqGTP is the highly efficient PLCβ activator, Gβγ also stimulates PLCβ and induces PIP2 hydrolysis^33^. We have shown that PIP2 hydrolysis induced by Gβγ alone is modest compared to the robust hydrolysis induced by GαqGTP^34^. Since α2AR is a Gi/o-GPCR, upon activation, Gβγ should be the only available PLCβ activator. Therefore, we examined whether depletion of plasma membrane-bound PIP2 by Gβγ retards PIP3 generation. We coexpressed α2AR, GRPR (Gq-GPCR), mCh-PH (PIP2 sensor), and Akt-PH-Venus (PIP3 sensor) in RAW 264.7 cells. Upon addition of 100 μM norepinephrine and 1 μM bombesin simultaneously, both α2AR and GRPR were activated, and PIP2 hydrolysis was observed (Fig. 2A). Interestingly, despite the robust PIP2 hydrolysis, we also observed a simultaneous PIP3 production and its subsequent attenuation. This showed that, even with significantly reduced PIP2 availability at the plasma membrane (Fig. 2A, PM-bound PIP2 and PIP3 plot), PI3K could still generate PIP3. We also examined PIP2 dynamics at the plasma membrane during Gi/o-GPCR-induced PIP3 level adapts. We coexpressed the same DNA combination as above and activated only α2AR. PIP2 sensor fluorescence at the plasma membrane remained unchanged during the PIP3 generation and its adaptation (Fig. 2B). These two conditions, in the presence (Fig. 2A) and absence (Fig. 2B) of Gq-GPCR activity, have created two distinct PIP2 availabilities at the plasma membrane for PI3Ks to generate PIP3. The PIP3 generation was significantly slower (∼2.5 fold) in the presence of Gq-GPCR activation compared to that in the absence of Gq-GPCR activation (0.0111 vs. 0.02858 s^−1^) (Fig. 2C - one-way ANOVA: *F*_1, 33_ = 47.05388, *p* = 7.87 × 10^−8^, Supplementary Table S2 A and B). This can be explained assuming the distinct concentrations of PIP2 available for PI3K, where PIP3 generation is impeded under low PIP2 abundance. However, the PIP3 adaptation rates or extents with and without Gq-GPCR activation showed no significant difference (adaptation rate: one-way ANOVA: *F*_1, 33_ = 2.36795, *p* = 0.133, adaptation extent: one-way ANOVA: *F*_1, 33_ = 0.32997, *p* = 0.56957) (Fig. 2C, Supplementary Table S3 A and B, Supplementary Table S4 A and B). This data suggests that while PIP2 availability at the plasma membrane could affect the magnitude and kinetics of PIP3 generation, it does not influence PIP3 adaptation. There have been discussions about the specificity of the PIP2 sensor and its ability to interact with PIP3^15^. Nevertheless, our data also show that even with strong PIP3 production at the plasma membrane, the PIP2 sensor stayed cytosolic after PIP2 hydrolysis (Fig. 2A, 100 and 400 s, and plots). This clearly shows that under the experimental conditions we employed, the PIP2 sensor is sufficiently specific for PIP2.

**Figure 2.**
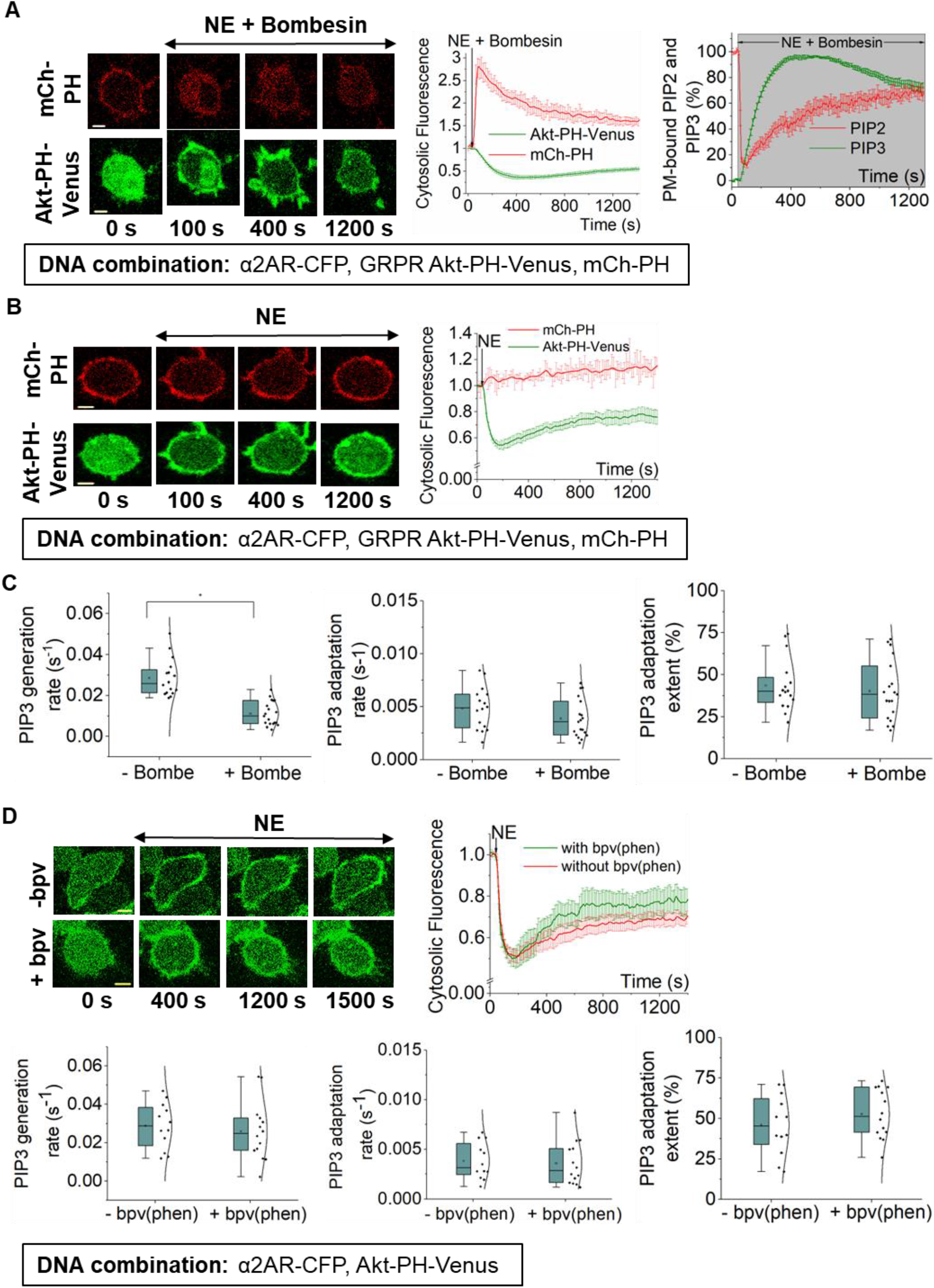
PIP3 adaptation is not a result of substrate (PIP2) depletion. **(A)** RAW264.7 cells expressing α2AR-CFP, GRPR, Akt-PH-Venus, and mCh-PH exhibited concurrent PIP2 hydrolysis - PIP3 production, and PIP3 adaptation upon simultaneous GRPR and α2AR activation at 1 min using 100 μM NE and 1 μM bombesin, respectively. Cells were imaged using 515 nm (for PIP3 sensor), and 594 nm (for PIP2 sensor). The corresponding plot shows the dynamics of PIP2 hydrolysis (red) and PIP3 production and adaptation (green). The plot compares the normalized PIP2 (red) and PIP3 (green) sensor dynamics at the PM. The two plots show that the changes in the sensor fluorescence in cytosol or the PM can be used to capture the dynamics of PIP2 and PIP3 at the PM (n = 19). **(B)** RAW264.7 cells expressing α2AR-CFP, GRPR, Akt-PH-Venus, and mCh-PH exhibited PIP3 production and attenuation upon α2AR activation (at 1 min), while PIP2 at the PM stayed intact. The corresponding plot shows the dynamics of PIP3 (green) and PIP2 (red) (n = 16). **(C)** The whisker box plots show PIP3 generation rates, adaptation rates, and adaptation extents for A and B experiments indicating the influence of PIP2 hydrolysis on PIP3. PIP3 generation rate in cells with PIP2 hydrolysis (+Bombesin) was significantly lower compared to cells without PIP2 hydrolysis (-Bombesin). However, the rate and extent of PIP3 adaptation under both conditions showed no significant difference. **(D)** RAW264.7 cells expressing α2AR-CFP and Akt-PH-Venus exhibited PIP3 production and attenuation upon α2AR activation in the presence or absence of phosphatase inhibitor bpV(phen) (5 μM). The corresponding curve plot shows the dynamics of PIP3 production and adaptation in the presence of bpV(phen) (green) (n=14), and in the absence of bpV(phen) (red) (n=12). The whisker box plots show PIP3 generation rates, adaptation rates, and adaptation extents in the presence and absence of bpV(phen). The generation and adaptation rates and adaptation extents showed no significant difference under both conditions. Statistical comparisons were performed using One-way-ANOVA (p < 0.05). Average curves were plotted using cells from ≥ 3 independent experiments. ‘n’ denotes the number of cells’ data used to plot the average curve. The error bars represent SEM. Scale bar = 5 μm. PIP2: phosphatidylinositol 4,5-bisphosphate; GRPR: gastrin-releasing peptide receptor

Cellular phosphatases such as PTEN reduce cellular PIP3 levels and regulate downstream signaling^35-38^. However, PTEN is not the only PIP3 phosphatase in cells. RNA seq data of RAW264.7 cells show a significantly higher expression of phosphatases, including PTEN, Inpp5d, and Inpp5b (Supplementary Fig. S3). Therefore, to examine whether phosphatases regulate PIP3 adaption, we employed the phosphatase inhibitor, bpV(phen), which has been shown to inhibit most phosphatases, including the above^39,40^. Cells expressing α2AR and Akt-PH-Venus were pre-incubated with 5 μM inhibitor for 30 minutes, and norepinephrine was added to activate α2AR. Upon activation, both control and phosphatase-inhibited cells showed PIP3 production and subsequent adaptation (Fig. 2D). The PIP3 generation rates (one-way ANOVA: *F*_1, 24_ = 0.29921, *p* = 0.58943, Supplementary Table S5 A and B), PIP3 adaptation rates (one-way ANOVA: *F*_1, 24_ = 0.08259, *p* = 0.77629, Supplementary Table S6 A and B), and the PIP3 adaptation extents (one-way ANOVA: *F*_1, 24_ = 1.04804, *p* = 0.31617, Supplementary Table S7 A and B) do not show a significant difference. This showed that neither PIP2 depletion nor phosphatase activity influenced PIP3 adaptation.

### 2.3 PIP3 generated by insulin receptors does not show adaptation

PIP3 is produced upon activating distinct signaling pathways, including GPCRs and Receptor Tyrosine Kinases (RTKs). RTKs induce PIP3 production by activating PI3Kα and β family members^41^, independent of G proteins. Since the PIP3 adaptation could occur due to upstream or downstream signaling of PIP3, similar to GPCR-induced, we examined whether the PIP3 generated upon RTK pathway activation is also susceptible to adaptation. Insulin receptors (InsR) are RTKs and induce PIP3 generation upon activation by insulin^17^. If GPCRs or G proteins play a significant role in PIP3 adaptation, we expect insulin-mediated PIP3 to be adaptation-resistant. We selected CHO cells since they express endogenous InsR^42^, As a control, we first examined whether CHO cells also show an adapting PIP3 response to Gi/o-GPCR-G protein activation. We expressed α2AR and Akt-PH-Venus in CHO cells. Upon addition of 100 μM norepinephrine, cells exhibited PIP3 generation (Fig. 3A, 400 and 800 s), followed by its usual adaptation seen in RAW 264.7 cells (Fig. 3A, 1600 s). We next monitored InsR-induced PIP3 generation upon addition of 10 μg/ml insulin. Though cells showed a robust PIP3 generation upon insulin addition, unlike the GPCR-induced, PIP3 remained intact after receptor activation (Fig. 3B, 1200 s). This data clearly indicated that the PIP3 adaptation is not due to negative feedback from RTK signaling.

**Figure 3.**
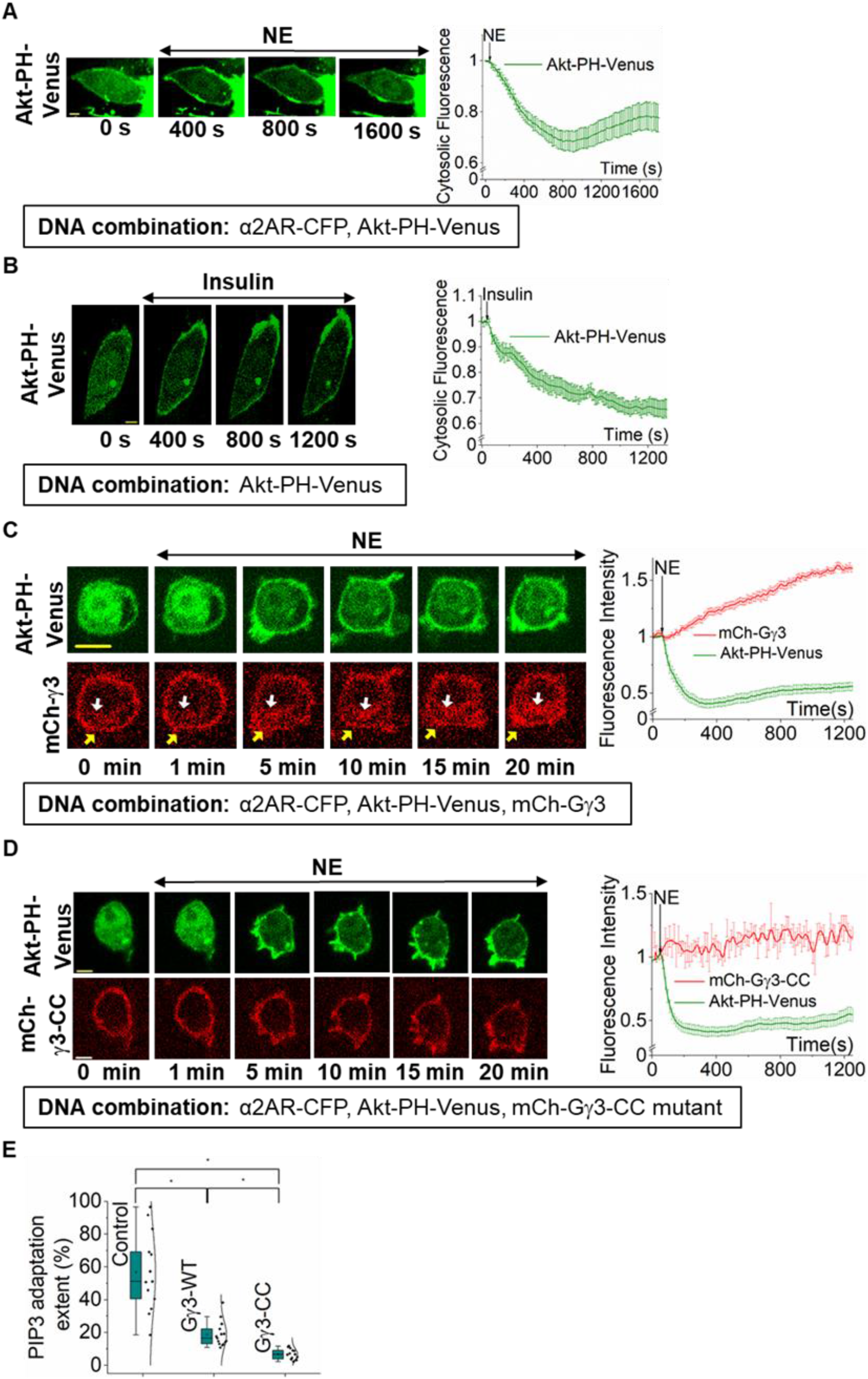
Gβγ and PIP3 adaptation. **(A)** CHO cells expressing α2AR-CFP and Akt-PH-Venus exhibited a robust PIP3 production and a subsequent adaptation upon α2AR activation with 100 μM NE. Sensor recruitment and reversal to and from the PM show PIP3 production and its partial adaptation, respectively. The corresponding plot shows dynamics of PIP3 production and adaptation (n = 8). **(B)** CHO cells expressing Akt-PH-Venus exhibited adaptation-resistant PIP3 response upon activation of endogenous insulin receptors (InsR). Insulin (10 μg/ml) was added at 1 min to activate InsR. Though cells showed a gradual PIP3 generation after insulin addition, contrary to the GPCR-induced, the adaptation of the produced PIP3 was not observed. The corresponding plot shows dynamics of PIP3 with InsR activation (n = 13). **(C)** RAW264.7 cells expressing α2AR-CFP, Akt-PH-Venus and mCh-Gγ3 exhibited simultaneous mCh-Gγ3 translocation and PIP3 production upon α2AR activation (at 1 min). The translocated mCherry-Gγ3 stayed at IMs (white arrows), while PIP3 level reached an equilibrium after adaptation. In addition to 515 nm excitation for YFP, 594 nm excitation was used to capture mCherry. The corresponding plot shows mCh-Gγ3 (red) and PIP3 (green) dynamics in the cytosol of the cells (n = 20). **(D)** RAW264.7 cells expressing α2AR-CFP, Akt-PH-Venus and mCh-Gγ3-CC mutant exhibited PIP3 production upon α2AR activation. As expected, the images and the plot show that translocation deficient Gγ mutant (Gγ3-CC) remained at the PM despite receptor activation. PIP3 showed no significant adaptation (n = 16). **(E)** The whisker box plots show the extents of % PIP3 adaptation in RAW264.7 cells with endogenous Gβγ (Control), or Gγ3 WT, or Gγ3-CC mutant. Extent of PIP3 adaptation was quantified using the increase in the mean cytosolic fluorescence due to PIP3 adaptation. The observed % adaptations were significantly different from each other (p < 0.05). Averages were plotted using cells from n≥ 3 independent experiments. ‘n’ denotes the number of cells’ data used to plot the average curve. The error bars represent SEM (standard error of mean). The scale bar = 5 μm. IMs: Internal membranes

### 2.4 Plasma membrane residency of Gβγ and the kinetics of the PIP3 adaptation

The glycerolipid group of Phosphatidylinositol is plasma membrane-bound, and therefore, to phosphorylate the substrate PIP2, PI3K subunits are recruited to the plasma membrane by plasma membrane-bound Gβγ^12^. Thus, Gβγ concentration at the plasma membrane should be a critical determinant of PIP3 production. Previously, it has been shown that Gβγ subunits generated upon GPCR activation at the plasma membrane reversibly translocate to internal membranes (IMs) in a Gγ-dependent manner^43,44^, and high membrane affinity Gγ types are translocation deficient^45^. Our previous work shows that PIP3 production upon GPCR activation requires the Gβγ composed of high membrane-affinity Gγ types^3^. Gγ3 shows the highest membrane affinity out of the 12 subtypes. Its expression facilitates the PIP3 production in HeLa cells, which typically do not show a significant PIP3 production upon GPCR activation due to the lack of high membrane-affinity Gγ expression at the endogenous level^28^. High membrane affinity Gβγ have the advantage of primarily residing at the plasma membrane even after being released from the heterotrimer^28^. Therefore, we hypothesized that the plasma membrane residency of Gβγ subunits and their translocation away from the plasma membrane regulate the observed PIP3 response adaptation. To examine whether Gβγ translocation and PIP3 response adaptation is related, we employed RAW 264.7 cells coexpressing α2AR, Akt-PH-Venus, and mCh-Gγ3. Activation of α2AR induced a robust PIP3 generation (Fig. 3C, green plot). As expected, Gγ3 showed slow and steady translocation indicated by mCherry fluorescence loss at the plasma membrane and gradual increase in the cell interior (Fig. 3C, red plot). Next, we analyzed the dynamics of Gγ3 translocation and PIP3 generation—adaptation. At the 5-minute mark, the majority of Gγ3 (∼ 89.18 ± 3.41 %) still stayed plasma membrane-bound (Fig. 3C-yellow arrows, Supplementary Fig. S1, and Supplementary Table S1), while PIP3 generation reached the maximum (Fig. 3C, 5 min, and plot). Even at the 20-minute mark, ∼ 65.039 ± 2.68 % of Gγ3 stayed plasma membrane-bound (Supplementary Fig. S1 and Supplementary Table S1), while PIP3 reduction has reached a steady state. Considering Gβγ is responsible for PIP3 generation, this data suggested that the percent PIP3 that remains at the plasma membrane is proportional to the Gβγ concentration. The extent of PIP3 response attenuation in WT-RAW264.7 cells (56.801 ± 6.61%) is ∼3 fold higher than that of the cells expressing Gγ3 (18.725 ± 1.93%) (Fig. 3E, Supplementary Table S8-A).

RNA seq data shows that RAW264.7 cells prominently express four major Gγ types at endogenous conditions, Gγ2 (36%), Gγ5 (14%), Gγ9 (17%), and Gγ12 (27%) (Supplementary Fig. S2 and Supplementary Table S9). Assuming that mRNA levels are proportional to protein expression, we can estimate the Gβγ concentration at the plasma membrane after its translocation reached the equilibrium in WT RAW264.7 cells since we have previously shown that Gγ2, 5, 9, and 12 translocate respectively 1.5, 3.8, 45, and 3.4 times faster than Gγ3^28^. For Gγ3-expressing cells, we assumed that the majority of Gγ (∼90%) in Gβγ is Gγ3. Since we did not express either Gα or Gβ, we also assumed nearly similar heterotrimer concentrations in both cellular conditions. To examine whether expressing high membrane-affinity Gγ increases the plasma membrane-bound Gβγ at the steady state, we compared percent plasma membrane-bound Gβγ3 and Gβγ9 at the steady state of PIP3 adaptation (∼20 mins). Here, our data showed that Gγ3-expressing cells have ∼6-fold higher Gβγ concentration at the plasma membrane compared to that of the Gγ9-expressing cells (Supplementary Fig. S1 and Supplementary Table S1). Since expression of Gγ3 increases the plasma membrane-bound Gβγ concentration at the steady state, we compared the PIP3 reduction extent between Gγ3-expressing cells and endogenous Gβγ-expressing cells. Here, endogenous cells showed ∼3-fold higher extent of PIP3 reduction (Fig. 3E- Control and Gγ3-WT, Supplementary Table S8 A and B). To further examine whether the plasma membrane residency of Gβγ is linked to the PIP3 reduction, we employed a translocation-deficient Gγ3 mutant (Gγ3-CC). This mutant has an additional *Cys* residue at the C-terminus, resulting in double lipidation, significantly enhancing its membrane affinity beyond Gγ3-WT^34^. We have also shown that Gγ3-CC encodes for a functional Gγ^34^. We coexpressed α2AR, Akt-PH-Venus, and mCh-Gγ3-CC mutant in RAW 264.7 cells. Activation of α2AR induced PIP3 generation; however, the Gγ3-CC mutant showed no translocation (Fig. 3D). Mutant-expressing cells showed a nearly 3-fold decrease in the extent of PIP3 adaptation (6.759 ± 0.85%) compared to Gγ3-WT-expressing cells (one-way ANOVA: *F*_1, 33_ = 56.87, *p* = 1.13 × 10^−8^) (Fig. 3E, Supplementary Table S8 A and B). Therefore, this data suggests that the continuous presence of active Gβγ on the plasma membrane in the presence of Gγ3-CC (Fig. 3D, red plot and cell images) allows for sustained PIP3 at the plasma membrane while preventing the PIP3 adaptation process (Fig. 3D, green plot and cell images). Further, membrane affinity differences in the above-considered Gγ types strongly suggest that the Gβγ loss from the plasma membrane due to translocation plays a crucial role in the dynamic adaptation of GPCR activation-induced PIP3.

### 2.5 Injection of Gβγ rescues adapted PIP3 generation induced by Gi/o pathway activation

We next examined whether the adapted PIP3 production can be rescued by injecting Gβγ into the system by activating Gq heterotrimers. RNAseq data shows that the endogenous Gαq expression in RAW264.7 cells is significantly lower compared to other Gα types, including Gαi/o (Supplementary Fig. S4 and Supplementary Table S11). Thus, it produces limited Gβγ upon Gq-GPCR activation, resulting in little to no PIP3 generation^46^. Since Gβγ is shared between different Gα types in a cell, we hypothesized that by expressing Gαq in RAW264.7 cells, we could increase Gq-GPCR activation-mediated Gβγ generation. We expressed GRPR (a Gq-coupled GPCR), Gαq-CFP, YFP-β1, and Lyn-HTH in RAW 264.7 cells. Since PLCβ-mediated PIP2 hydrolysis significantly impedes PIP3 production (Fig. 2A and C, Supplementary Table S2 A and B), we employed plasma membrane-targeted HTH (Helix-Turn-Helix) domain of PLCβ3 (from Y847-E884) since it competes with PLCβ to interact with GαqGTP, and has been shown to inhibit PIP2 hydrolysis^47^. The data show that Lyn-HTH completely inhibited Gαq-induced PLCβ stimulation; however, it did not prevent Gβγ translocation, indicating unperturbed heterotrimer activation (Fig. 4A).

**Figure 4.**
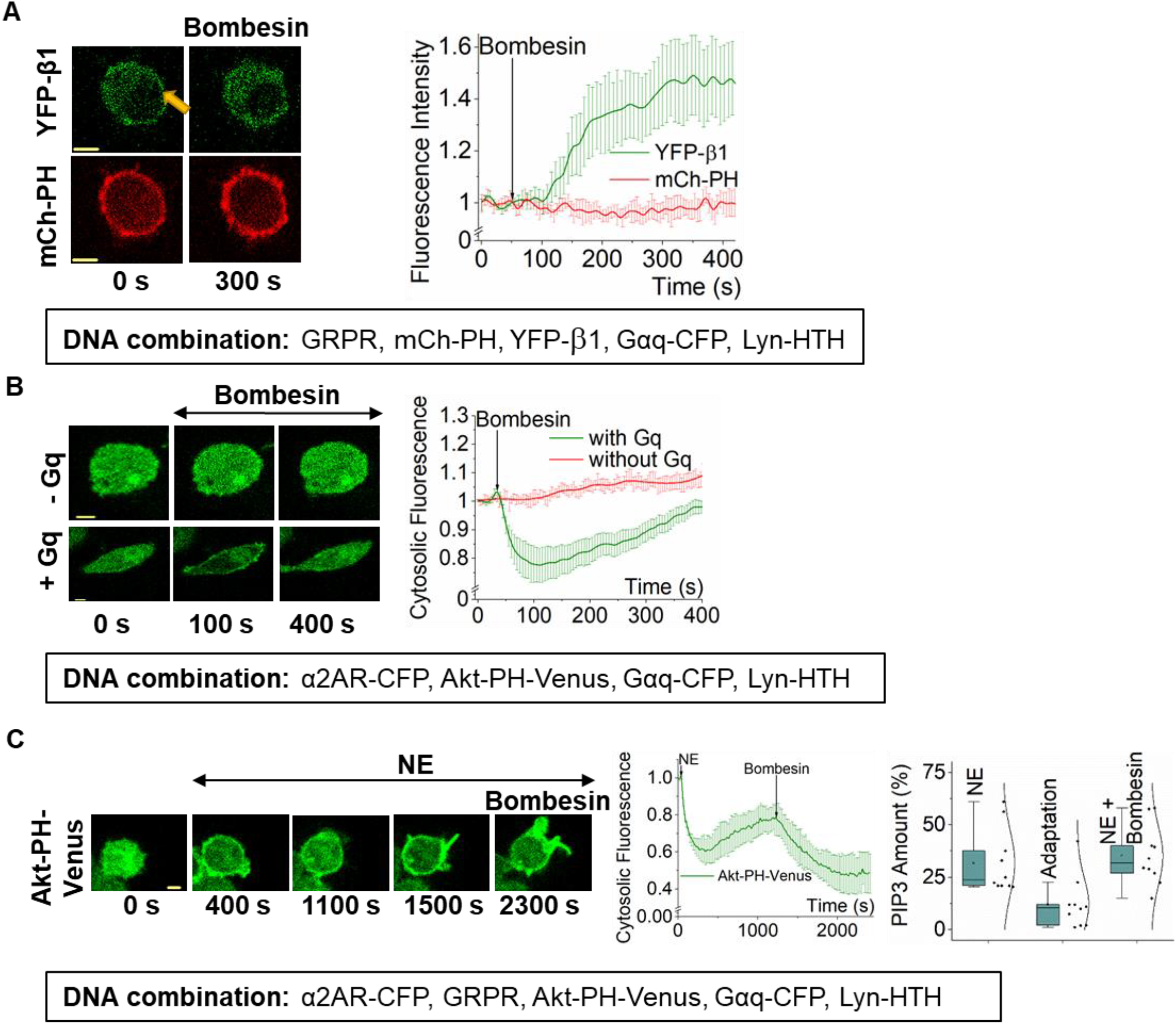
Despite the source, Gβγ entry induces PIP3 generation and loss results in adaptation. **(A)** RAW264.7 cells expressing GRPR, YFP-β1, mCh-PH, Gαq-CFP, and Lyn-HTH exhibited robust β1 translocation upon GRPR activation with bombesin (1 μM at 1 min) (n = 8). However, GRPR activation did not induce PIP2 hydrolysis due to the presence of Lyn-HTH. The plot shows the dynamics of β1 translocation (green) and PIP2 (red) in cells. **(B)** RAW264.7 cells expressing GRPR, Akt-PH-Venus, Gαq-CFP, and Lyn-HTH exhibited PIP3 production and adaptation upon GRPR activation with 1 μM bombesin (n = 8). However, control cells expressing only GRPR and Akt-PH-Venus did not show PIP3 production upon GRPR activation (n= 9). The plot shows PIP3 production and adaptation only in Gαq-expressing cells (green), however not in control cell lacking introduced Gαq. **(C)** RAW264.7 cells expressing α2AR-CFP, GRPR, Akt-PH-Venus, Gαq-CFP and Lyn-HTH exhibited PIP3 production and adaptation upon α2AR activation. After PIP3 adaptation, 1 μM bombesin was added (at 20 min) to activate GRPR. GRPR activation in these Gαq cells caused disruption of adaptation in the form of PIP3 increase. The plot shows the initial PIP3 generation and adaptation after α2AR activation, and subsequent GRPR-induced PIP3 regeneration in Gαq background (n = 10). Average curves were plotted using cells from ≥ 3 independent experiments. ‘n’ denotes the number of cells’ data used to plot the average curve. The error bars represent SEM (standard error of mean). The scale bar = 5 μm.

To examine Gq-GPCR activation-mediated Gβγ can generate PIP3, we expressed GRPR (a Gq-coupled GPCR), Gαq-CFP, Akt-PH-Venus, and Lyn-HTH in RAW 264.7 cells. As expected, a robust PIP3 generation and subsequent adaptation were observed in Gq-expressing cells upon GRPR activation (Fig. 4B, +Gq). However, in the absence of Gαq expression, GRPR activation did not induce PIP3 production (Fig. 4B, -Gq). This data also ruled out the possibility of Gαi exclusive requirement in PIP3 generation or its adaptation. Since GRPR activation in Gαq expressing cells exhibited a robust Gβγ generation that is even sufficient to induce a significant PIP3 generation, we next examined whether the PIP3 adaptation observed after Gi/o-GPCR could be rescued by injecting Gβγ using Gq-GPCR pathway activation. We expressed α2AR, GRPR, Gαq-CFP, Akt-PH-Venus, and Lyn-HTH in RAW 264.7 cells. We first activated α2AR and observed PIP3 generation (Fig. 4C, 400 s, and plots). We continued imaging cells for 20 minutes till PIP3 at the plasma membrane was reduced to a constant level indicating the maximum partial adaptation (Fig. 4C, 1100 s, and plots). We then activated Gq-coupled GRPR using 1 μM bombesin. The partially adapted PIP3 levels in cells were increased to the pre-adaptation level (compare Fig. 4C, 400 s, and 1500 s). This data suggests that Gβγ injection can rescue the PIP3 adaptation, further suggesting that the loss of plasma membrane-bound Gβγ maybe the molecular reason behind PIP3 adaptation. Considering the lack of PIP3 adaptation observed in translocation-incompetent Gγ3 and translocation-deficient Gγ3 CC-mutant cells, this PIP3 rescue observed upon Gβγ injection clearly indicates the Gβγ involvement in the PIP3 adaptation.

### 2.6 Localized optogenetic inhibition of active Gαi can disrupt the PIP3 adaptation process

Localized optical activation of a Gi/o-coupled GPCR, blue opsin, shows a sustained and adaptation-resistant localized PIP3 generation at the leading edge in migrating RAW264.7 cells (Fig. 5A- Localized)^48^. Here we used cells expressing blue opsin-mTurquoise and Akt-PH-mCh. After the addition of 1 μM 11-*cis*-retinal (to generate optically activable blue opsin), and while imaging the cells for mCherry, we exposed a confined plasma membrane region of a cell to 445 nm blue light by exposing an adjacent area to rectangular-shaped blue light pulse delivered at 1Hz. The opsin activation induced cell migration towards the localized blue light, which was steered to manage the migration direction. The cells showed a sustained localized PIP3. When a cell was exposed to blue light globally, although the PIP3 generation was observed, the PIP3 response adapted quickly (Fig. 5A- Global). Since the asymmetric GPCR activation-mediated PIP3 production was adaptation resistant, we examined whether breaking the signaling symmetry in a PIP3-adapted cell could also recover the cell from the PIP3 adaptation. Here we employed an optogenetic GTPase engineered using a truncated version of Regulator of G protein signaling 4 domain (RGS4Δ) that has been used to optically inhibit G protein activity in cells^49^. RGS4Δ domain accelerates GTP hydrolysis on Gαi/oGTP, sequestering Gβγ to form Gαβγ heterotrimers^50^. RGS4Δ is tethered to a cryptochrome 2-based CRY2-mCh-RGS4Δ. Upon blue light stimulation, CRY2 dimerizes with its plasma membrane-targeted binding partner, a truncated version of cryptochrome-interacting basic-helix-loop-helix (CIBN)^49^. We expressed α2AR-CFP, CRY2-mCh-RGS4Δ, CIBN-CAAX, and Akt-PH-Venus in RAW264.7 cells. While imaging cells for mCherry and Venus, we exposed a localized membrane region of the cell to blue light (Fig. 5B, blue box). The localized blue light recruited CRY2-mCh-RGS4Δ (Fig. 5B, 1 min). We then activated α2AR globally by adding 100 μM norepinephrine. Cells produced PIP3 only at the opposite side from the CRY2-mCh-RGS4Δ localization (Fig. 5B, 3 and 10 min). Here, the recruitment of RGS4Δ should significantly decrease the lifetime of GαGTP and Gβγ, reducing their concentration to a negligible level at the RGS4Δ localized side of the cell. However, on the opposite side, heterotrimers are activated, Gαi-GTP and Gβγ are generated, where Gβγ stimulates PI3Kγ to induce localized PIP3, orchestrating directional cell migration. Similar to the blue opsin induced (Fig. 5A- Localized), the continuous blue light-directed localized RGS4Δ resulted in adaptation-resistant PIP3 at the opposite edge of the cell (Fig. 5B). Upon termination of blue light, the localized PIP3 gradually disappeared (Fig. 5B, 15 min).

Next, in a cell expressing the same construct combination, we activated α2AR globally first and allowed PIP3 to be produced (Fig. 5C, 3 min) and adapted (Fig. 5C, 10 min). We then recruited CRY2-mCh-RGS4Δ to one side using localized blue light (Fig. 5C, 15 min, and 20 min). Localized recruitment of RGS4Δ disrupted the PIP3 response adaptation, and the opposite side of the cell showed PIP3 generation. Upon switching CRY2-mCh-RGS4Δ to the opposite side, we were able to switch the side of the PIP3 generation in the same cell (Fig. 5C, 25 min, and 30 min). Here, the data collectively show that, when the GPCR activation is global, the PIP3 adaptation is rapid, while localized GPCR activation, as well as asymmetric G protein activation, deliver sustained and localized PIP3 generations at the GPCR/G protein active site. Data also show that even in a cell with PIP3 adaptation incurred, introducing asymmetry to G protein activation breaks the PIP3 adaptation. Here, we utilized three distinct methods to induce asymmetric signaling. In the first two conditions, signaling asymmetry was introduced before the PIP3 adaptation, i. e. (i) localized GPCR activation through blue light-induced blue opsin activation, where both GPCRs and G proteins remained active, and (ii) global GPCR activation by adding norepinephrine to activate α2AR in a cell with one side of the plasma membrane of the cell with the inhibitor that eliminated activated G proteins. In both conditions, cells showed adaptation-resistant PIP3 generation. In the third condition, i. e. a cell with GPCR activated globally, and PIP3 response is adapted, G protein activity termination in one side of the plasma membrane rescued the cells from the adaptation. Collectively, this data suggests the involvement of G proteins, likely Gβγ, in the PIP3 adaptation mechanism.

**Figure 5.**
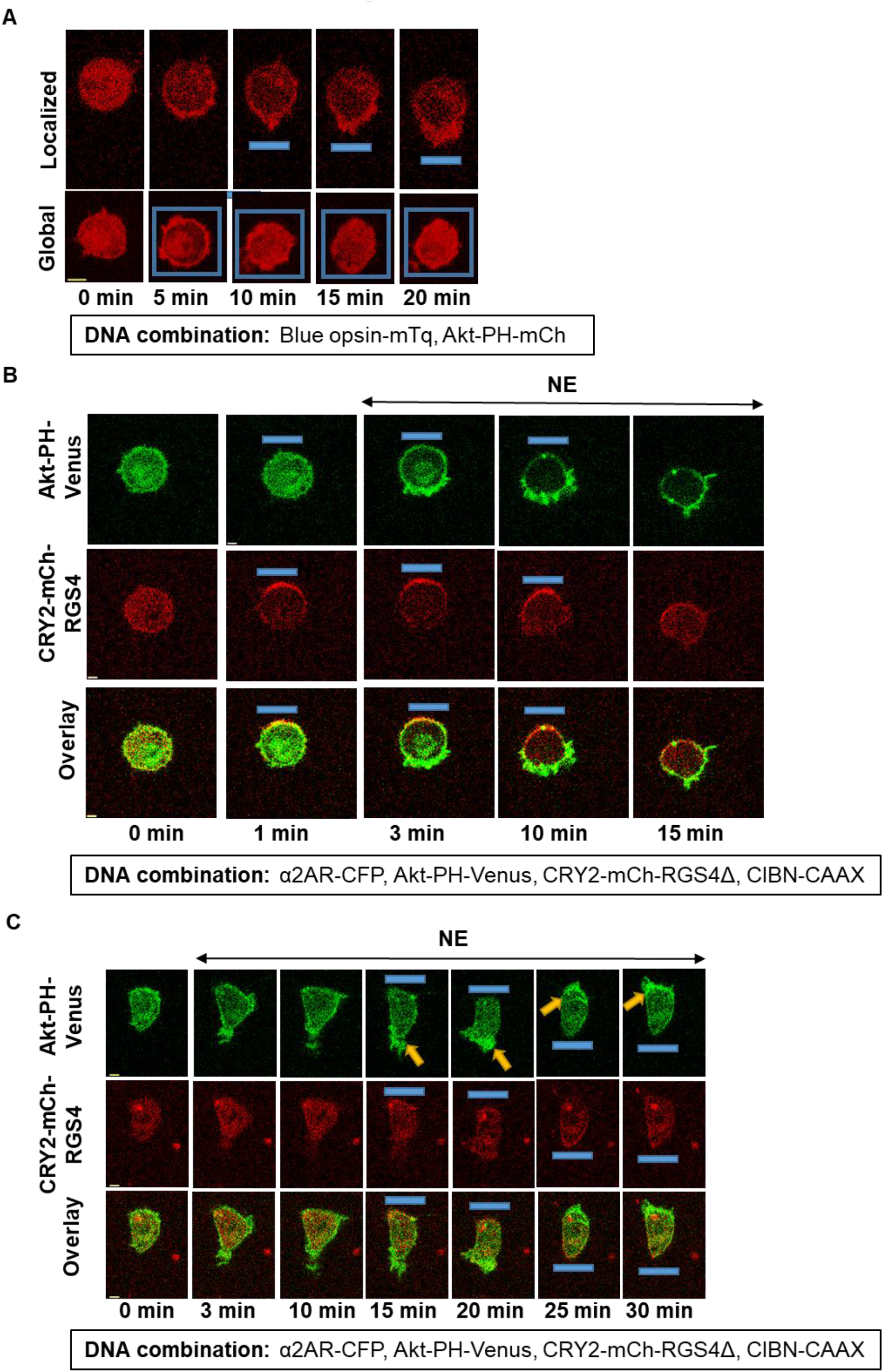
PIP3 generation upon localized GPCR-G protein activation is adaptation-resistant. **(A)** RAW264.7 cells expressing blue opsin-mTq and Akt-PH-mCh showed adaptation-resistant PIP3 upon localized activation, while global activation induced fast attenuating PIP3. Before activation cells were incubated with 1 μM 11-cis-retinal. Spatially confined blue light pulses (blue boxes) (1 Hz) was used activate by blue light locally and globally, while confocal imaging of mCherry using 594 nm excitation. Cells activated with localized blue light produced non-adapting PIP3 response. Cells also showed a directional migration towards the blue light. On the contrary, cells exposed to blue light globally initially showed a robust global PIP3 production. However, PIP3 in these cells showed the adaptation, similar to the adaptation observed upon activation of α2AR. **(B)** Localized acceleration of GTP on Gα in α2AR activated RAW264.7 cells also showed adaptation-resistant PIP3 generation. Cells expressing α2AR-CFP, Akt-PH-Venus, CRY2-mCh-RGS4Δ, and CIBN-CAAX showed localized PIP3 production when α2AR was activated in a cells after CRY2-mCh-RGS4Δ localized to one side of the cell. A localized blue light pulse was provided to recruit CRY2-mCh-RGS4Δ to one side of the cell. Once RGS4Δ is recruited, (indicated by mCherry fluorescence concentrating to one side of the cell), NE (100 μM) was added to the medium. This resulted in localized PIP3 at the opposite side to the blue pulse (yellow arrow). Cells were imaged using 515 nm (to capture PIP3 sensor), and 594 nm (to capture RGS4Δ). Although images show data from only one cell, experiments were conducted in multiple cells with >3 independent experiments to test the reproducibility of the results. **(C)** PIP3-adapted (post-GPCR activation) RAW264.7 cells show disruption of adaptation upon localized inhibition G protein activity. Cells expressed α2AR-CFP, Akt-PH-Venus, CRY2-mCh-RGS4Δ, and CIBN-CAAX. Initially α2AR was activated with 100 μM NE, and PIP3 was produced and significantly adapted in 10 mins. Upon blue light induced recruitment of CRY2-mCh-RGS4Δ to one side of the cell, PIP3 was produced at the opposite side (yellow arrow, 15 and 20 min), indicating adaptation disruption. When the opposite side of the same cell was exposed to blue light (yellow arrow, 25 and 30 min), PIP3 was produced at the opposite side of localized RGS4Δ. Experiments were conducted in multiple cells with >3 independent experiments to test the reproducibility of the results. The blue box indicates the blue light. The scale bar = 5 μm. mTq: mTurquoise; FRAPPA: fluorescence recovery after photo-bleaching and photo-activation; RGS4: Regulator of G protein signaling 4

### 2.7 Sustained localized PIP3 production is facilitated by heterotrimer shuttling to the GPCR active site

Next, we examined molecular reasoning for localized G protein activation to deliver adaptation-resistant PIP3 generation. Since RAW264.7 cells are much smaller than most other cell types, we used HeLa cells to visualize G protein movement between subcellular compartments in global and localized signaling activations.

First, we examined G protein redistribution in a cell where localized RGS4Δ actively reduces the concentration of active G proteins (GαGTP and Gβγ) in one side of a cell in which Gi/o-GPCRs are activated globally. We expressed α2AR, YFP-Gβ1, CRY2-mCh-RGS4Δ, and CIBN-CAAX in HeLa cells. YFP-Gβ1 initially showed a plasma membrane distribution, indicating that it is in the Gαβγ heterotrimer (Fig. 6A, 0 s, and plot). Activation of α2AR induced a gradual Gβ1 translocation to internal membranes (Fig. 6A, 250 s, and plot). Next, we recruited CRY2-mCh-RGS4Δ to one side of the cell using localized blue light (blue box). Gβ1 partially recovered to the RGS4Δ-recruited side upon blue light exposure (Fig. 6A 400 s, white arrow, and green plot). However, there was no Gβ1 recovery at the opposite of CRY2-mCh-RGS4Δ-recruited side (Fig. 6A 400 s, grey arrow, and black plot). In a separate experiment, we activated α2AR with 100 μM norepinephrine, and upon Gβ1 translocation (Fig. 6B, 350 s, and plot), we recruited CRY2-mCh-RGS4Δ using global blue light. Demonstrating the GTPase activity of RGS4Δ, cells showed reverse translocation of Gβ1 to the plasma membrane (Fig. 6B, 600 s, yellow arrow and plot). We then kept the cell in the dark for 5 minutes. Not only CRY2-mCh-RGS4Δ fully returned to the cytosol, we also observed endomembrane localized Gβ1, indicating Gβγ translocation (Fig. 6B, 1000 s, and plot). These observations suggest that when RGS4Δ is recruited to the plasma membrane locally, heterotrimers should be formed at the recruited side, while the heterotrimer concentration reaches near zero on the opposite side. Therefore, upon localized RGS4Δ recruitment, the cell should have a heterotrimer concentration gradient across the cell. In a concentration gradient, molecules move from higher to lower until an equilibrium is achieved. Therefore, heterotrimers from the RGS4Δ side should shuttle to the opposite side. Since the opposite side heterotrimer concentration is near zero, shuttling should continue indefinitely, as long as global GPCR and local RGS4Δ activities are maintained.

**Figure 6.**
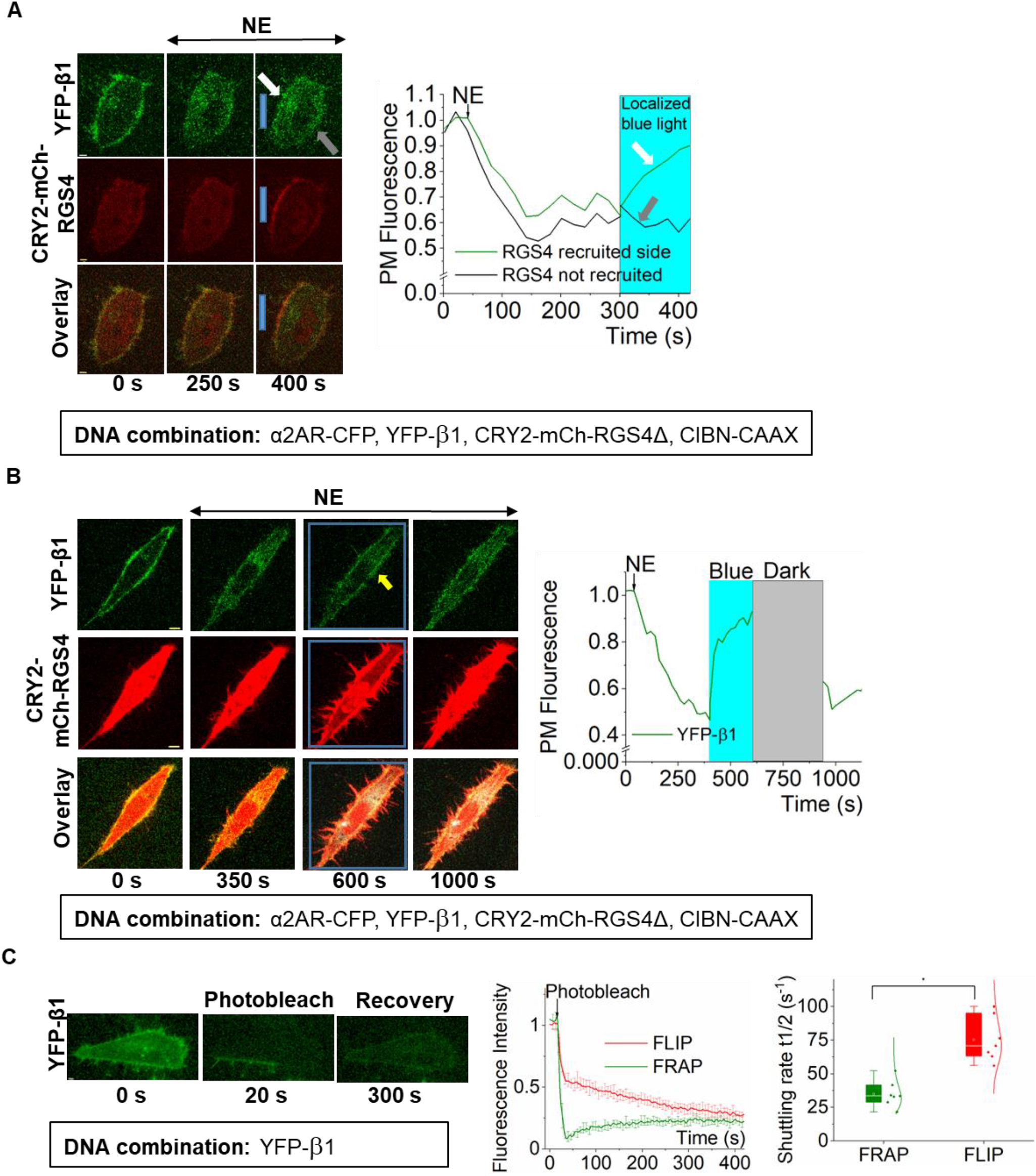
Heterotrimer concentration gradient across the cell breaks PIP3 adaptation. **(A)** Optical targeting of RGS4Δ to on side of a cell induces localized heterotrimer regeneration. HeLa cells expressing α2AR-CFP, YFP-Gβ1, CRY2-mCh-RGS4Δ, and CIBN-CAAX initially exhibited robust Gβ1 translocation upon α2AR activation (250 s). Then, upon localized blue light-induced recruitment of CRY2-mCh-RGS4Δ resulted in reverse translocation of Gβ1 from IMs to PM, indicated by the increase in PM-bound YFP fluorescence (white arrow, green plot), suggesting heterotrimer regeneration. No increase was observed at the opposite side (grey arrow, black plot), suggesting heterotrimers are still dissociated. The corresponding plot shows the dynamics of YFP-β1 with localized inhibition of CRY2-mCh-RGS4Δ. **(B)** Reversible blue light targeting of RGS4Δ to the PM allows optogenetic control of heterotrimer concentration at the PM. Upon α2AR activation, cells showed YFP-Gβ1 translocation (350 s). Blue light exposure recruited CRY2-mCh-RGS4Δ to the PM, which induced reverse translocation of Gβ1 to the PM (600 s). Termination of blue light that resulted in the release of PM-bound CRY2-mCh-RGS4Δ again resulted in YFP-Gβ1 translocation (1000 s). Thus ON-OFF blue light allowed reversible control of heterotrimer concentration. The corresponding plot shows the dynamics of YFP-β1 with global inhibition by CRY2-mCh-RGS4Δ. **(C)** Fluorescence recovery after subcellular photobleaching (FRAP) in HeLa cells expressing YFP-Gβ1. The corresponding plot shows fluorescence recovery after photobleaching (FRAP) in photobleached PM regions of the cell (green), and fluorescence loss after photobleaching (FLIP) in non-photobleached PM regions. The whisker plot shows the shuttling half times Gβ1 in FRAP (green) and FLIP (red) (n = 7). Average curves were plotted using cells from ≥ 3 independent experiments. ‘n’ denotes the number of cells’ data used to plot the average curve. The error bars represent SEM (standard error of mean). The scale bar = 5 μm.

To examine how fast heterotrimers shuttle from one side to the other in a living cell, we expressed YFP-Gβ1 in HeLa cells and photobleached YFP except for a small plasma membrane strip (Fig. 6C, 20 s). We then examined the shuttling of YFP-Gβ1 to the opposite bleached area by calculating Fluorescence loss in photobleaching (FLIP) and Fluorescence recovery after photobleaching (FRAP) at the opposite sides. The shuttling half time of heterotrimers in FRAP was 34.99 ± 3.66 s, and FLIP was 75.38 ± 6.23 s. When the time duration from RGS4Δ recruitment to the initiation of the PIP3 generation at the opposite side (∼5 mins) is considered, the heterotrimer shuttling rate is relatively minor. Therefore, it is likely that heterotrimers generated at the RGS4Δ side (as in Figs. 5C, 6A, and 6B) reach the opposite side, and significantly elevate Gβγ concentration compared to the Gβγ concentration in globally GPCR activated cells (without RGS4Δ recruitment). We propose that this elevated localized Gβγ in asymmetrically G protein activated cells allow for adaptation-resistant PIP3 generation. In a globally GPCR activated cell, at the equilibrium after GPCR activation, heterotrimer concentration at the plasma membrane should go to near zero. Though the Gβγ concentration should go up at the plasma membrane initially, upon translocation, it should be significantly reduced. Since PIP3 generation is Gβγ-dependent, over time PIP3 level at the plasma membrane should significantly reduce, or in other words, adapted, due to the reduced production. As we showed above (Fig. 5B), in a globally GPCR activated cell with RGS4Δ recruited, the free Gβγ is constantly sequestered in the heterotrimer due to RGS4Δ-induced efficient GαGDP formation, resulting in no PIP3 production.

## 3. Conclusion

Cells and tissues develop adaptation mechanisms to avoid overstimulation despite continuous stimuli. Gi-coupled GPCR activation induces a fast-attenuating PIP3 response. Our results indicate that PIP3 attenuation is not due to GPCR desensitization, PIP2 loss at the plasma membrane, PIP3 degradation by cellular phosphatases, or negative feedback from downstream effectors of PIP3. Here we show that the attenuation of the GPCR-induced PIP3 response is indeed an adaptation since the attenuation incurs while continuous stimulation (both the ligand and functional receptors). Here, we identified Gβγ translocation from the plasma membrane to endomembranes that control their concentration at the plasma membrane inner leaflet as the underlying mechanisms for the observed PIP3 adaptation (Fig. 7). Since Gβγ recruits PI3Kγ to the plasma membrane where the substrate, PIP2, resides^51^, our data show that plasma membrane-residing-ability or lack thereof of Gβγ regulates PI3K activity—PIP3 generation. Plasma membrane-residence of Gβγ is linked to their membrane affinity, which is governed by the associated Gγ subtype^28^. Our findings further demonstrate that this Gβγ-regulated PIP3 adaptation is Gγ type-dependent. We also decoded the unexpected, adaptation-resistant PIP3 response triggered by the asymmetric GPCR or G protein activation. We showed that the excess heterotrimer availability and the elevated G protein activation at the localized plasma membrane-area, a scenario that is absent when GPCRs-G proteins are globally activated, signals for the sustained localized PIP3. We previously showed that such localized PIP3 sets the migration direction, as well as likely to be involved in neuronal symmetry breaking and differentiation^52^. Further, the ability of cells to produce rapidly adapting PIP3 upon global GPCR activation should therefore act as a signal for cells to cease the cell-fate-deciding behaviors, including migration, creating autonomous regulation of crucial cell behaviors possible.

**Figure 7.**
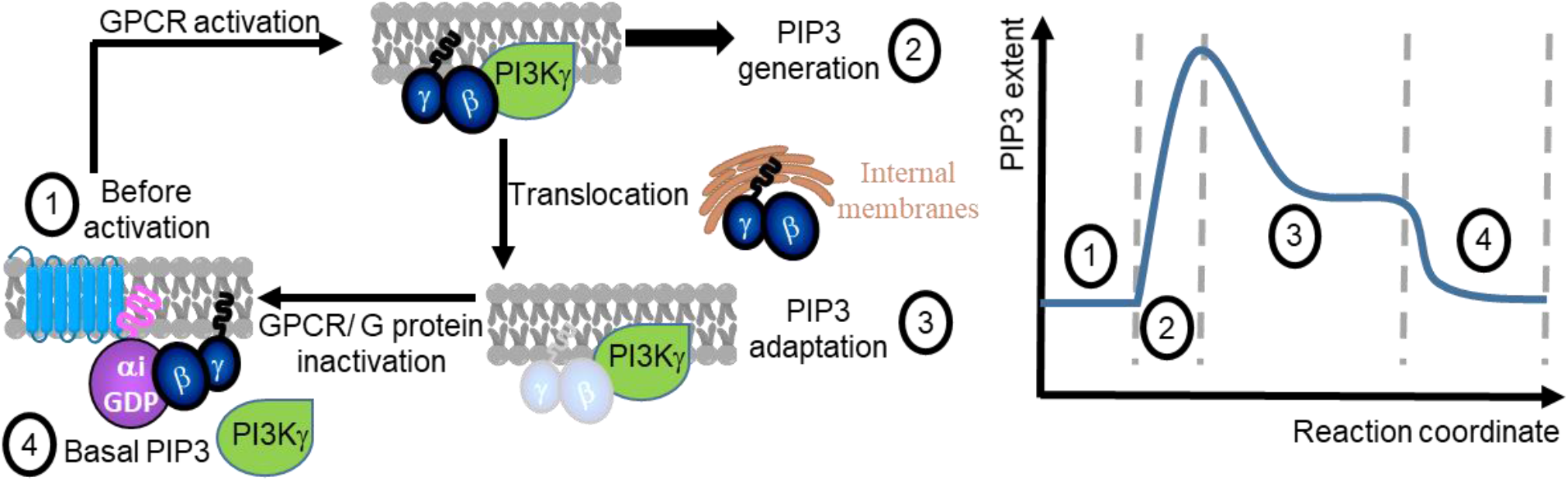
Proposed mechanisms of GPCR-G protein-induced PIP3 regulation. Gi/o-coupled GPCR activation induces robust free Gbg, which stimulates PI3Kg resulting in PIP3 generation (1). The subsequent partial PIP3 adaptation (2) is due to the translocation of Gbg away rom the plasma membrane to the internal membranes. The steady state of partial PIP3 adaptation is achieved at the equilibrium of Gbg translocation between plasma membrane and the internal membranes. Complete inactivation of GPCR or G proteins result in PIP3 level returning to the basal level.

Gγ shows distinct cell and tissue type-specific distribution patterns.^53^ Therefore, our findings allow for cells with low plasma membrane-affinity Gβγ to show faster and greater PIP3 adaptation while setting the molecular machinery for cells expressing high plasma membrane-affinity Gβγ to have slow-adapting PIP3 responses. Therefore, our newly found regulatory mechanisms in GPCR-G protein-PI3K-PIP3 signaling can allow knowledge-based predictions in signaling on outcomes of different cells and tissues. Since GPCR-G protein signaling is one of the major drug targets and PI3Kγ-PIP3 signaling is implicated in oncogenesis^9,54,55^, our findings will help realize the abundance, impact, and enormity of distinct and diverse cell-tissue-specific signaling regimes of the same pathway, and their implications in health and disease.

## 4. Materials and methods

### 4.1 Reagents

The reagents used were as follows; Norepinephrine (NE) (Sigma Aldrich), Yohimbine, Bombesin (Tocris Bioscience), PBS, Insulin (Sigma corporation, St Louis, MO), 11-*cis*-retinal (National Eye Institute). Stock solutions of compounds were prepared according to manufacturers’ recommendations. Before adding to cells, all stock solutions were diluted in 1% Hank’s balanced salt solution (HBSS) or regular cell culture medium.

### 4.2 DNA constructs and cell lines

DNA constructs used were as follows; DNA constructs used for Blue opsin-mTurquoise, mCherry-Gγ3-CC mutant, fluorescently tagged Akt-PH, CRY2-mCherry-RGS4Δ, and PH, Lyn-HTH have been described previously^34,45,52,56,57^. CIBN-CAAX was cloned into the pcDNA3.1 vector in our lab. NES-Venus-mini-Gi was kindly provided by Professor N. Lambert’s laboratory, Augusta University, Augusta, GA. GRPR was a kind gift from the laboratory of Dr. Zhou-Feng Chen at Washington University, St Louis, MO. Fluorescently tagged Gγ3 and Gγ9 subunits, YFP-β1, and αq–CFP were kindly provided by Professor N. Gautam’s laboratory, Washington University, St Louis, MO. All constructs were cloned by Gibson assembly cloning (NEB). Cloned cDNA constructs were confirmed by sequencing (Genewiz). Cell lines used were as follows: RAW264.7, CHO, and HeLa cells were purchased from the American Tissue Culture Collection (ATCC).

### 4.3 Cell culture and transfections

RAW264.7 cells were cultured in Roswell Park Memorial Institute (RPMI) 1640 medium (Corning, Manassas, VA) supplemented with 10% heat-inactivated dialyzed fetal bovine serum (DFBS, Atlanta Biologicals, GA) and 1% penicillin-streptomycin (PS, 10,000 U/ml stock) and grown at 37 °C with 5% CO_2_. CHO cells were maintained in Dulbecco’s modified Eagle medium: Nutrient Mixture F-12 (DMEM/F-12, Cellgro) with 10% DFBS and 1% PS. HeLa cells were cultured in minimum essential medium (Cellgro) with 10% DFBS and 1% PS. All cells were cultured in 35 mm, 60 mm, or 100 mm cell culture dishes (Celltreat). DNA transfections were performed using either Lipofectamine®2000 reagent (for CHO and HeLa cells) or electroporation (for RAW264.7 cells). For lipofection, cells were seeded onto 35 mm cell culture-grade glass-bottomed dishes (Cellvis) at a density of 8 × 104 cells and were transfected, the next day, with appropriate DNA constructs using Lipofectamine®2000 reagent (Invitrogen). The transfection media was changed to regular cell culture media after 5 hours and the cells were imaged 16 hours post-transfection. For electroporation of RAW264.7 cells, the electroporation solution was prepared with the Nucleofector solution (82 μL), Supplement solution (18 μL), and appropriate volumes of DNA constructs. For each experiment, ∼2-4 million cells were electroporated using the T020 method of the Nucleofector™ 2b device (Lonza). Immediately after electroporation, cells were mixed with cell culture medium at 37 °C and seeded onto 35 mm cell culture-grade glass-bottomed dishes coated with poly-L-lysine. Cells were imaged ∼5-6 hours post-electroporation.

### 4.4 Live cell imaging, image analysis, and data processing

The methods, protocols, and parameters for live-cell imaging are adapted from previously published work^44,58,59^. Briefly, live-cell imaging experiments were performed using a spinning disk (Yokogawa CSU-X1, 5000 rpm) XD confocal TIRF imaging system composed of a Nikon Ti-R/B inverted microscope with a 60X, 1.4 NA oil objective and iXon ULTRA 897BVback-illuminated deep-cooled EMCCD camera. Photoactivation and Spatio-temporally controlled light exposure on cells in regions of interest (ROI) were performed using a laser combiner with 40-100 mW solid-state lasers (445, 488, 515, and 594 nm) equipped with Andor® FRAP-PA unit (fluorescence recovery after photobleaching and photoactivation), controlled by Andor iQ 3.1 software (Andor Technologies, Belfast, United Kingdom). Fluorescent sensors such as Akt-PH-mCherry, mCherry-γ3, mCherry-γ9, mCherry-PH, and CRY2-mCherry-RGS4Δ were imaged using 594 nm excitation− 624 nm emission settings; Akt-PH-Venus and Venus-mini-Gi were imaged using 515 nm excitation and 542 nm emission; Gαq-CFP was imaged using 445 nm excitation and 478 nm emission. For global and confined optical activation of CRY2 expressing cells, the power of 445 nm solid-state laser was adjusted to 5 mW. Additional adjustments of laser power with 0.1%-1% transmittance were achieved using Acousto-optic tunable filters (AOTF). Ophir PD300-UV light meter was used for laser power measurements. Data acquisition, time-lapse image analysis, processing, and statistical analysis were performed as explained previously^58^. Briefly, Time-lapse images were analyzed using Andor iQ 3.1 software by acquiring the mean pixel fluorescence intensity changes of the entire cell or the selected area/regions of interest (ROIs).

### 4.5 Statistical data analysis

All experiments were repeated multiple times to test the reproducibility of the results. Statistical analysis and data plot generation were done using OriginPro software (OriginLab®). Results were analyzed from multiple cells and represented as mean ± SEM. The exact number of cells used in the analysis is given in respective figure legends. PIP3 generation and adaptation rates were calculated using the Nonlinear Curve Fitting tool (NLFit) in OriginPro. Each plot was fitted to DoseResp (Dose-Response) function under the Pharmacology category in OriginPro. The mean values of hill slopes (P) obtained for each nonlinear curve fitting are presented as mean rates of PIP3 generation or adaptation. PIP3 dynamics plots fit to the Michaelis-Menten function were used to determine the t_½_of PIP3 generation and adaptation. The obtained mean values of Km were taken as the mean t_½_One-way ANOVA statistical tests were performed using OriginPro to determine the statistical significance between two or more populations of signaling responses. Tukey’s mean comparison test was performed at the p < 0.05 significance level for the one-way ANOVA statistical test.

## Acknowledgment

We acknowledge Dr. N. Gautam for providing us with plasmid DNA of various G protein subunits and for the RNA seq data. We thank the National Eye Institute for providing 11-*cis*-retinal. We also thank Senuri Piyawardana, Kiran Ghotra, Hasheena Rajapakshage, and Chathuri Rajarathna for their comments.

## Author contributions

D.W. and K.R. conducted many of the experiments and performed the data analysis. S.U. conducted the Gγ3 translocation and PIP3 generation experiments. D.K. performed mini Gi recruitment with PIP3 generation-adaptation experiment. M.T. assisted in insulin receptor-induced PIP3 generation experiments. D.W. performed the statistical analysis. A.K., K.R., and D.W. conceptualized the project and wrote the manuscript.

## References

1 Alarcón, C. d. l. H., Pennadam, S. & Alexander, C. Stimuli responsive polymers for biomedical applications. Chemical Society Reviews 34, 276–285, doi:10.1039/B406727D (2005).

2 Janetopoulos, C., Jin, T. & Devreotes, P. Receptor-Mediated Activation of Heterotrimeric G-Proteins in Living Cells. Science 291, 2408–2411, doi:10.1126/science.1055835 (2001).

3 Hoeller, O., Gong, D. & Weiner, O. D. How to understand and outwit adaptation. Dev Cell 28, 607–616, doi:10.1016/j.devcel.2014.03.009 (2014).

4 Ferrell, J. E. Perfect and Near-Perfect Adaptation in Cell Signaling. Cell Systems 2, 62-67, doi:https://doi.org/10.1016/j.cels.2016.02.006 (2016).

5 Stauffer, T. P., Ahn, S. & Meyer, T. Receptor-induced transient reduction in plasma membrane PtdIns(4,5)P2 concentration monitored in living cells. Current Biology 8, 343–346, doi:10.1016/s0960-9822(98)70135-6 (1998).

6 Bononi, A. et al. Protein kinases and phosphatases in the control of cell fate. Enzyme research 2011, 329098, doi:10.4061/2011/329098 (2011).

7 Carnero, A. & Paramio, J. M. The PTEN/PI3K/AKT Pathway in vivo, Cancer Mouse Models. Frontiers in Oncology 4, 252 (2014).

8 Vanhaesebroeck, B., Guillermet-Guibert, J., Graupera, M. & Bilanges, B. The emerging mechanisms of isoform-specific PI3K signalling. Nature Reviews Molecular Cell Biology 11, 329–341, doi:10.1038/nrm2882 (2010).

9 Jean, S. & Kiger, A. A. Classes of phosphoinositide 3-kinases at a glance. Journal of Cell Science 127, 923–928, doi:10.1242/jcs.093773 (2014).

10 Jean, S. & Kiger, A. A. Classes of phosphoinositide 3-kinases at a glance. Journal of Cell Science 127, 923–928, doi:10.1242/jcs.093773 (2014).

11 Brock, C. et al. Roles of Gβγ in membrane recruitment and activation of p110γ/p101 phosphoinositide 3-kinase γ. Journal of Cell Biology 160, 89–99, doi:10.1083/jcb.200210115 (2003).

12 Vadas, O. et al. Molecular determinants of PI3Kγ-mediated activation downstream of G-protein– coupled receptors (GPCRs). Proceedings of the National Academy of Sciences 110, 18862–18867, doi:10.1073/pnas.1304801110 (2013).

13 Dbouk, H. A. et al. G Protein Coupled Receptor Mediated Activation of p110beta by Gbetagamm Is required for Cellular Transformation and Invasiveness. Science Signaling 5, ra89–ra89, doi:doi:10.1126/scisignal.2003264 (2012).

14 Hemmings, B. A. & Restuccia, D. F. Pi3k-pkb/akt pathway. Cold Spring Harbor perspectives in biology 4, a011189 (2012).

15 Czech, M. P. PIP2 and PIP3: Complex Roles at the Cell Surface. Cell 100, 603–606, doi:10.1016/S0092-8674(00)80696-0 (2000).

16 Riehle, R. D., Cornea, S. & Degterev, A. Role of phosphatidylinositol 3,4,5-trisphosphate in cell signaling. Advances in experimental medicine and biology 991, 105–139, doi:10.1007/978-94-007-6331-9_7 (2013).

17 Shepherd, P. R., Withers, D. J. & Siddle, K. Phosphoinositide 3-kinase: the key switch mechanism in insulin signalling. Biochemical Journal 333, 471–490, doi:10.1042/bj3330471 (1998).

18 Hassan, B., Akcakanat, A., Holder, A. M. & Meric-Bernstam, F. Targeting the PI3-kinase/Akt/mTOR signaling pathway. Surg Oncol Clin N Am 22, 641–664, doi:10.1016/j.soc.2013.06.008 (2013).

19 Zhang, M., Goswami, M. & Hereld, D. Constitutively Active G Protein-coupled Receptor Mutants Block Dictyostelium Development. Molecular Biology of the Cell 16, 562–572, doi:10.1091/mbc.e04-06-0456 (2005).

20 Zhang, J., Feng, H., Xu, S. & Feng, P. Hijacking GPCRs by viral pathogens and tumor. Biochemical pharmacology 114, 69–81, doi:10.1016/j.bcp.2016.03.021 (2016).

21 O’Neill, P. R., Gautam, N. & Haastert, P. V. Subcellular optogenetic inhibition of G proteins generates signaling gradients and cell migration. Molecular Biology of the Cell 25, 2305–2314, doi:10.1091/mbc.e14-04-0870 (2014).

22 Janetopoulos, C., Ma, L., Devreotes, P. N. & Iglesias, P. A. Chemoattractant-induced phosphatidylinositol 3,4,5-trisphosphate accumulation is spatially amplified and adapts, independent of the actin cytoskeleton. Proceedings of the National Academy of Sciences 101, 8951–8956, doi:10.1073/pnas.0402152101 (2004).

23 Meili, R. et al. Chemoattractant-mediated transient activation and membrane localization of Akt/PKB is required for efficient chemotaxis to cAMP in Dictyostelium. EMBO J 18, 2092–2105, doi:10.1093/emboj/18.8.2092 (1999).

24 Parent, C. A., Blacklock, B. J., Froehlich, W. M., Murphy, D. B. & Devreotes, P. N. G Protein Signaling Events Are Activated at the Leading Edge of Chemotactic Cells. Cell 95, 81-91, doi:https://doi.org/10.1016/S0092-8674(00)81784-5 (1998).

25 Servant, G. et al. Polarization of chemoattractant receptor signaling during neutrophil chemotaxis. Science 287, 1037–1040, doi:10.1126/science.287.5455.1037 (2000).

26 Barkai, N. & Leibler, S. Robustness in simple biochemical networks. Nature 387, 913–917, doi:10.1038/43199 (1997).

27 Weis, R. M. & Koshland, D. E., Jr. Reversible receptor methylation is essential for normal chemotaxis of Escherichia coli in gradients of aspartic acid. Proceedings of the National Academy of Sciences 85, 83–87, doi:10.1073/pnas.85.1.83 (1988).

28 Senarath, K. et al. Gγ identity dictates efficacy of Gβγ signaling and macrophage migration.. Journal of Biological Chemistry 293, 2974–2989, doi:10.1074/jbc.RA117.000872 (2018).

29 Kelly, E., Bailey, C. P. & Henderson, G. Agonist-selective mechanisms of GPCR desensitization. Br J Pharmacol 153 Suppl 1, S379–S388, doi:10.1038/sj.bjp.0707604 (2008).

30 Black, J. B., Premont, R. T. & Daaka, Y. Feedback regulation of G protein-coupled receptor signaling by GRKs and arrestins. Semin Cell Dev Biol 50, 95–104, doi:10.1016/j.semcdb.2015.12.015 (2016).

31 Wan, Q. et al. Mini G protein probes for active G protein-coupled receptors (GPCRs) in live cells. Journal of Biological Chemistry, doi:10.1074/jbc.RA118.001975 (2018).

32 Koyasu, S. The role of PI3K in immune cells. Nature Immunology 4, 313–319, doi:10.1038/ni0403-313 (2003).

33 Boyer, J. L., Graber, S. G., Waldo, G. L., Harden, T. K. & Garrison, J. C. Selective activation of phospholipase C by recombinant G-protein alpha- and beta gamma-subunits. Journal of Biological Chemistry 269, 2814-2819, doi:https://doi.org/10.1016/S0021-9258(17)42015-1 (1994).

34 Kankanamge, D. et al. Dissociation of the G protein betagamma from the Gq-PLCbeta complex partially attenuates PIP2 hydrolysis. Journal of Biological Chemistry 296, doi:10.1016/j.jbc.2021.100702 (2021).

35 Leslie, N. R., Batty, I. H., Maccario, H., Davidson, L. & Downes, C. P. Understanding PTEN regulation: PIP2, polarity and protein stability. Oncogene 27, 5464–5476, doi:10.1038/onc.2008.243 (2008).

36 Vazquez, F. et al. Tumor suppressor PTEN acts through dynamic interaction with the plasma membrane. Proceedings of the National Academy of Sciences 103, 3633–3638, doi:10.1073/pnas.0510570103 (2006).

37 Myers, M. P. et al. The lipid phosphatase activity of PTEN is critical for its tumor supressor function. Proceedings of the National Academy of Sciences 95, 13513–13518 (1998).

38 Maehama, T. & Dixon, J. E. The tumor suppressor, PTEN/MMAC1, dephosphorylates the lipid second messenger, phosphatidylinositol 3,4,5-trisphosphate. Journal of Biological Chemistry 273, 13375–13378, doi:10.1074/jbc.273.22.13375 (1998).

39 Schmid, A. C., Byrne, R. D., Vilar, R. & Woscholski, R. Bisperoxovanadium compounds are potent PTEN inhibitors. FEBS letters 566, 35–38, doi:10.1016/j.febslet.2004.03.102 (2004).

40 Batty, I. H. et al. The control of phosphatidylinositol 3,4-bisphosphate concentrations by activation of the Src homology 2 domain containing inositol polyphosphate 5-phosphatase 2, SHIP2. Biochemical journal 407, 255–266, doi:10.1042/BJ20070558 (2007).

41 Molinaro, A. et al. Insulin-Driven PI3K-AKT Signaling in the Hepatocyte Is Mediated by Redundant PI3Kα and PI3Kβ Activities and Is Promoted by RAS. cell metabolism 29, 1400-1409.e1405, doi:10.1016/j.cmet.2019.03.010 (2019).

42 Myers, M. G., Backer, J. M., Siddle, K. & White, M. F. The insulin receptor functions normally in Chinese hamster ovary cells after truncation of the C terminus. Journal of Biological Chemistry 266, 10616–10623 (1991).

43 Ajith Karunarathne, W. K., O’Neill, P. R., Martinez-Espinosa, P. L., Kalyanaraman, V. & Gautam, N. All G protein betagamma complexes are capable of translocation on receptor activation. Biochemical and biophysical research communications 421, 605–611, doi:10.1016/j.bbrc.2012.04.054 (2012).

44 Senarath, K., Ratnayake, K., Siripurapu, P., Payton, J. L. & Karunarathne, A. Reversible G Protein betagamma9 Distribution-Based Assay Reveals Molecular Underpinnings in Subcellular, Single-Cell, and Multicellular GPCR and G Protein Activity. Analytical Chemistry 88, 11450–11459, doi:10.1021/acs.analchem.6b02512 (2016).

45 Senarath, K. et al. Gγ identity dictates efficacy of Gβγ signaling and macrophage migration. Journal of Biological Chemistry 293, 2974–2989, doi:10.1074/jbc.RA117.000872 (2018).

46 Kankanamge, D. et al. G protein αq exerts expression level-dependent distinct signaling paradigms. Cellular Signalling 58, 34-43, doi:https://doi.org/10.1016/j.cellsig.2019.02.006 (2019).

47 Charpentier, T. H. et al. Potent and Selective Peptide-based Inhibition of the G Protein Gαq. Journal of Biological Chemistry 291, 25608–25616, doi:10.1074/jbc.M116.740407 (2016).

48 Siripurapu, P., Kankanamge, D., Ratnayake, K., Senarath, K. & Karunarathne, A. Two independent but synchronized Gbetagamma subunit-controlled pathways are essential for trailing-edge retraction during macrophage migration. Journal of Biological Chemistry 292, 17482–17495, doi:10.1074/jbc.M117.787838 (2017).

49 Kennedy, M. J. et al. Rapid blue-light-mediated induction of protein interactions in living cells. Nat Methods 7, 973–975, doi:10.1038/nmeth.1524 (2010).

50 Srinivasa, S. P., Bernstein, L. S., Blumer, K. J. & Linder, M. E. Plasma membrane localization is required for RGS4 function in Saccharomyces cerevisiae. Proceedings of the National Academy of Sciences 95, 5584–5589, doi:10.1073/pnas.95.10.5584 (1998).

51 Tannert, A., Voigt, P., Burgold, S., Tannert, S. & Schaefer, M. Signal amplification between Gbetagamma release and PI3Kgamma-mediated PI(3,4,5)P3 formation monitored by a fluorescent Gbetagamma biosensor protein and repetitive two component total internal reflection/fluorescence redistribution after photobleaching analysis. Biochemistry 47, 11239–11250, doi:10.1021/bi800596b (2008).

52 Karunarathne, W. K. A., Giri, L., Kalyanaraman, V. & Gautam, N. Optically triggering spatiotemporally confined GPCR activity in a cell and programming neurite initiation and extension. Proceedings of the National Academy of Sciences 110, E1565–E1574, doi:10.1073/pnas.1220697110 (2013).

53 Tennakoon, M. et al. Subtype-dependent regulation of Gβγ signalling. Cellular Signalling 82, 109947, doi:https://doi.org/10.1016/j.cellsig.2021.109947 (2021).

54 Zhao, L. & Vogt, P. K. Class I PI3K in oncogenic cellular transformation. Oncogene 27, 5486–5496, doi:10.1038/onc.2008.244 (2008).

55 Hauser, A. S., Attwood, M. M., Rask-Andersen, M., SchiöTh, H. B. & Gloriam, D. E. Trends in GPCR drug discovery: new agents, targets and indications. Nature Reviews Drug Discovery 16, 829–842, doi:10.1038/nrd.2017.178 (2017).

56 Chisari, M., Saini, D. K., Cho, J.-H., Kalyanaraman, V. & Gautam, N. G protein subunit dissociation and translocation regulate cellular response to receptor stimulation. PloS one 4, e7797 (2009).

57 O’Neill, P. R., Kalyanaraman, V. & Gautam, N. Subcellular optogenetic activation of Cdc42 controls local and distal signaling to drive immune cell migration. Molecular biology of the cell 27, 1442–1450 (2016).

58 Ratnayake, K., Payton, J. L., Lakmal, O. H. & Karunarathne, A. Blue light excited retinal intercepts cellular signaling. Scientific Reports 8, 10207, doi:10.1038/s41598-018-28254-8 (2018).

59 Ratnayake, K. et al. Blue light-triggered photochemistry and cytotoxicity of retinal. Cellular Signalling, 109547, doi:https://doi.org/10.1016/j.cellsig.2020.109547 (2020).

