## Supplementary Information for "Molecular regulation of GPCR-G-protein-governed PIP3 generation and its adaptation"

---

**Figure S1**

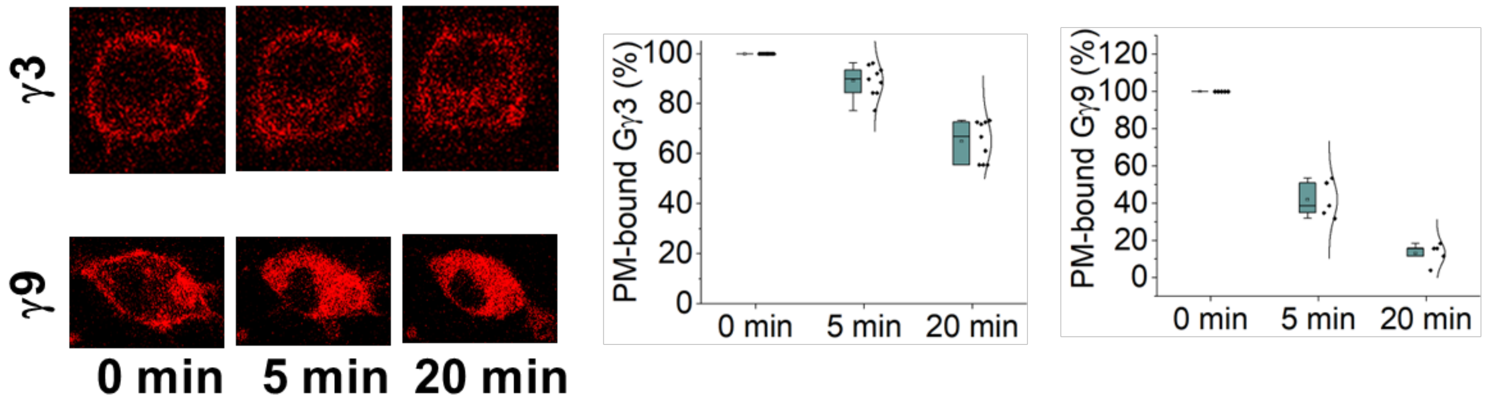


**Figure S1.** The extent of PM-bound Gγ3 and Gγ9 at 5 minutes and 20 minutes after Gi/o-coupled α2AR activation.

**Table S1: Plasma membrane-bound percent Gγ3 and Gγ9 at the steady state of PIP3 adaptation**

| Gγ type | PM-bound % Gγ (before activation) | PM-bound % Gγ 5 mins after activation | PM-bound % Gγ 20 mins after activation |
|---|---|---|---|
| Gγ3 | 100 (Overexpression) | 89.12048 | 65.03899 |
| Gγ9 | 100 (Overexpression) | 41.94931 | 13.11796 |

**Table S2-A: One-way ANOVA statistics for PIP3 generation rates with and without Gq-GPCR activation**

| **Descriptive Statistics- PIP3 production rates** | | | | |
|---|---|---|---|---|
| | N Analysis | Mean | Standard Deviation | SE of Mean |
| without bombesin | 16 | 0.02858 | 0.00914 | 0.00229 |
| with Bombesin | 19 | 0.0111 | 0.00581 | 0.00133 |

**Table S2-B**

| **Overall ANOVA- PIP3 production rates** | | | | | |
|---|---|---|---|---|---|
| | DF | Sum of Squares | Mean Square | F Value | Prob>F |
| Model | 1 | 0.00265 | 0.00265 | 47.05388 | 7.87283E-8 |
| Error | 33 | 0.00186 | 5.64147E-5 | | |
| Total | 34 | 0.00452 | | | |

At the 0.05 level, the population means are **significantly** different.

**Table S3-A: One-way ANOVA statistics for PIP3 adaptation rates with and without Gq-GPCR activation**

| **Descriptive Statistics- PIP3 adaptation rates** | | | | |
|---|---|---|---|---|
| | N Analysis | Mean | Standard Deviation | SE of Mean |
| Without bombesin | 16 | 0.00484 | 0.00199 | 4.98471E-4 |
| With bombesin | 19 | 0.00385 | 0.0018 | 4.13079E-4 |

**Table S3-B**

| **Overall ANOVA- PIP3 adaptation rates** | | | | | |
|---|---|---|---|---|---|
| | DF | Sum of Squares | Mean Square | F Value | Prob>F |
| Model | 1 | 8.46654E-6 | 8.46654E-6 | 2.36795 | 0.13338 |
| Error | 33 | 1.17991E-4 | 3.57547E-6 | | |
| Total | 34 | 1.26457E-4 | | | |

At the 0.05 level, the population means are **not significantly** different.

**Table S4-A: One-way ANOVA statistics for PIP3 adaptation extent with and without Gq-GPCR activation**

| **Descriptive Statistics- PIP3 adaptation extent** | | | | |
|---|---|---|---|---|
| | N Analysis | Mean | Standard Deviation | SE of Mean |
| Without Bombesin | 16 | 43.51647 | 15.62973 | 3.90743 |
| With bombesin | 19 | 40.24849 | 17.65828 | 4.05109 |

**Table S4-B**

| **Overall ANOVA- PIP3 adaptation extent** | | | | | |
|---|---|---|---|---|---|
| | DF | Sum of Squares | Mean Square | F Value | Prob>F |
| Model | 1 | 92.76091 | 92.76091 | 0.32997 | 0.56957 |
| Error | 33 | 9276.99677 | 281.12111 | | |
| Total | 34 | 9369.75768 | | | |

At the 0.05 level, the population means are **not significantly** different.

**Table S5-A: One-way ANOVA statistics for PIP3 generation rates with and without bpv(phen) inhibitor**

| **Descriptive Statistics- PIP3 generation rates** | | | | |
|---|---|---|---|---|
| | N Analysis | Mean | Standard Deviation | SE of Mean |
| Without bpv(phen) | 12 | 0.02884 | 0.01193 | 0.00344 |
| With bpv(phen) | 14 | 0.02589 | 0.01506 | 0.00403 |

**Table S5-B**

| **Overall ANOVA- PIP3 generation rates** | | | | | |
|---|---|---|---|---|---|
| | DF | Sum of Squares | Mean Square | F Value | Prob>F |
| Model | 1 | 5.62906E-5 | 5.62906E-5 | 0.29921 | 0.58943 |
| Error | 24 | 0.00452 | 1.8813E-4 | | |
| Total | 25 | 0.00457 | | | |

At the 0.05 level, the population means are **not significantly** different.

**Table S6-A: One-way ANOVA statistics for PIP3 adaptation rates with and without bpv(phen) inhibitor**

| **Descriptive Statistics- PIP3 adaptation rates** | | | | |
|---|---|---|---|---|
| | N Analysis | Mean | Standard Deviation | SE of Mean |
| Without bpv(phen) | 12 | 0.00382 | 0.00185 | 5.33617E-4 |
| With bpv(phen) | 14 | 0.00359 | 0.00222 | 5.94051E-4 |

**Table S6-B**

| **Overall ANOVA- PIP3 adaptation rates** | | | | | |
|---|---|---|---|---|---|
| | DF | Sum of Squares | Mean Square | F Value | Prob>F |
| Model | 1 | 3.5036E-7 | 3.5036E-7 | 0.08259 | 0.77629 |
| Error | 24 | 1.01814E-4 | 4.24225E-6 | | |
| Total | 25 | 1.02164E-4 | | | |

At the 0.05 level, the population means are **not significantly** different.

**Table S7-A: One-way ANOVA statistics for PIP3 adaptation extent with and without bpv(phen) inhibitor**

| **Descriptive Statistics- PIP3 adaptation extent** | | | | |
|---|---|---|---|---|
| | N Analysis | Mean | Standard Deviation | SE of Mean |
| Without bpv(phen) | 12 | 46.01196 | 18.5537 | 5.35599 |
| With bpv(phen) | 14 | 52.79968 | 15.26877 | 4.08075 |

**Table S7-B**

| **Overall ANOVA- PIP3 adaptation extent** | | | | | |
|---|---|---|---|---|---|
| | DF | Sum of Squares | Mean Square | F Value | Prob>F |
| Model | 1 | 297.70369 | 297.70369 | 1.04804 | 0.31617 |
| Error | 24 | 6817.39873 | 284.05828 | | |
| Total | 25 | 7115.10242 | | | |

At the 0.05 level, the population means are **not significantly** different.

**Table S8-A: One-way ANOVA statistics for PIP3 adaptation extent in Gγ3-CC mutant and Gγ3-WT expressing cells with control cells**

| **Descriptive Statistics- PIP3 adaptation extent** | | | | |
|---|---|---|---|---|
| | N Analysis | Mean | Standard Deviation | SE of Mean |
| Control | 13 | 56.80103 | 23.85578 | 6.6164 |
| Gγ3-WT | 15 | 18.72531 | 7.47989 | 1.9313 |
| Gγ3-CC | 14 | 6.75863 | 3.17492 | 0.84853 |

**Table S8-B**

| **Overall ANOVA- PIP3 adaptation extent** | | | | | |
|---|---|---|---|---|---|
| | DF | Sum of Squares | Mean Square | F Value | Prob>F |
| Model | 2 | 18298.74998 | 9149.37499 | 46.08066 | 5.35237E-11 |
| Error | 39 | 7743.4999 | 198.55128 | | |
| Total | 41 | 26042.24988 | | | |

At the 0.05 level, the population means are **significantly** different.

**Table S9: Normalized Gγ subtype expression profile in RAW 264.7 cells using RNA seq data**

| Gγ subtype | Expression (number of reads) | α4A-tubulin expression (number of reads) | Relative Expression (normalized to α4A-tubulin expression) | % expression |
|---|---|---|---|---|
| Gγ1 | 0 | 7404.33 | 0 | 0 |
| Gγ2 | 1444.33 | 7404.33 | 0.195066 | 35.90 |
| Gγ3 | 4.67 | 7404.33 | 0.00063 | 0.12 |
| Gγ4 | 1.67 | 7404.33 | 0.000225 | 0.04 |
| Gγ5 | 560.33 | 7404.33 | 0.075676 | 13.93 |
| Gγ7 | 42 | 7404.33 | 0.005672 | 1.04 |
| Gγ8 | 3.33 | 7404.33 | 0.00045 | 0.08 |
| Gγ9 | 684.67 | 7404.33 | 0.092468 | 17.02 |
| Gγ10 | 195.33 | 7404.33 | 0.026381 | 4.85 |
| Gγ11 | 3.67 | 7404.33 | 0.000495 | 0.09 |
| Gγ12 | 1082.67 | 7404.33 | 0.146221 | 26.91 |
| Gγ13 | 1 | 7404.33 | 0.000135 | 0.02 |

**Figure S2**

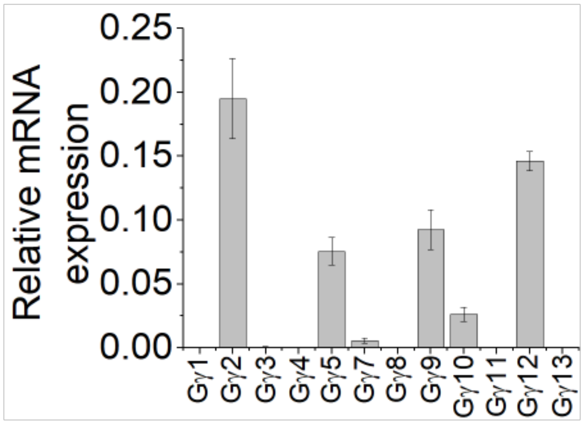


**Figure S2.** Relative mRNA expression levels of the 12 different Gγ subtypes in RAW264.7 cells. RAW264.7 cells show significant expression of Gγ2, 12, 9, and 5 subtypes. Values were normalized to α4A-tubulin.

**Table S10: Normalized expression profile of phosphatases in RAW 264.7 cells using RNA seq data**

| Phosphatase type | Expression (number of reads) | α4A-tubulin expression (number of reads) | Relative Expression (normalized to α4A-tubulin expression) |
|---|---|---|---|
| PTEN | 5484 | 7404.33 | 0.740647 |
| Inpp5a | 296.33 | 7404.33 | 0.040022 |
| Inpp5b | 1798 | 7404.33 | 0.242831 |
| Inpp5d | 6651 | 7404.33 | 0.898258 |
| Inpp5e | 617.33 | 7404.33 | 0.083375 |
| Inpp5f | 738.33 | 7404.33 | 0.099716 |
| Inpp5k | 408.67 | 7404.33 | 0.055193 |
| Inpp4a | 283 | 7404.33 | 0.038221 |

**Figure S3**

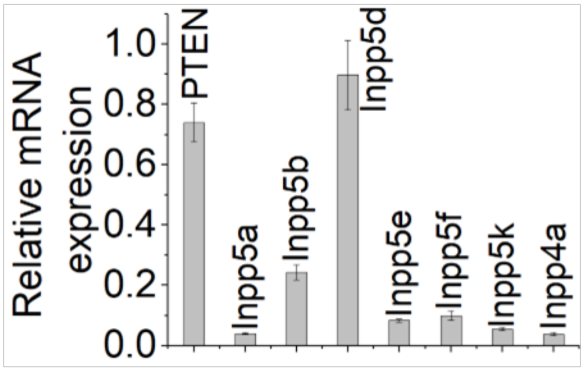


**Figure S3.** Relative mRNA expression levels of the PIP3 phosphatases in RAW264.7 cells. RAW264.7 cells show significant expression of PTEN and Inpp5d. Values were normalized to α4A-tubulin.

**Table S11: Normalized Gαi/o and Gαq/11 expression profile in RAW 264.7 cells using RNA seq data**

| Gα type | Expression (number of reads) | α4A-tubulin expression (number of reads) | Relative Expression (normalized to α4A-tubulin expression) |
|---|---|---|---|
| Gαi/o | 42384.67 | 7404.33 | 5.724306 |
| Gαq/11 | 3902.67 | 7404.33 | 0.527079 |

**Figure S4**

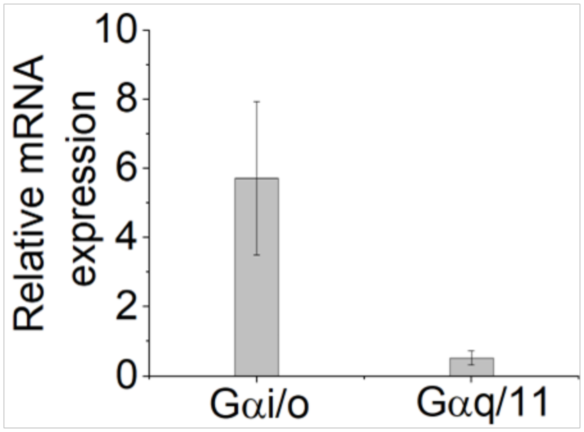


**Figure S4.** Relative mRNA expression levels of the Gαi/o and Gαq/11 in RAW264.7 cells. RAW264.7 cells show 10-fold higher expression of Gαi/o compared to Gαq/11. Values were normalized to α4A-tubulin.
